# A structural atlas of the developing zebrafish telencephalon based on spatially-restricted transgene expression

**DOI:** 10.1101/2022.03.02.482259

**Authors:** Katherine J. Turner, Thomas A. Hawkins, Pedro M. Henriques, Leonardo E. Valdivia, Isaac H. Bianco, Stephen W. Wilson, Mónica Folgueira

## Abstract

Zebrafish telencephalon acquires an everted morphology by a two-step process that occurs from 1 to 5 days post-fertilization (dpf). Little is known about how this process affects the positioning of discrete telencephalic cell populations, hindering our understanding of how eversion impacts telencephalic structural organisation. In this study, we characterise the neurochemistry, cycle state and morphology of an EGFP positive (+) cell population in the telencephalon of Et(*gata2*:*EGFP*)^bi105^ transgenic fish during eversion and up to 20dpf. We map the transgene insertion to the *early-growth-response-gene-3* (*egr3*) locus and show that EGFP expression recapitulates endogenous *egr3* expression throughout much of the pallial telencephalon. Using the *gata2:EGFP*^bi105^ transgene, in combination with other well-characterised transgenes and structural markers, we track the development of various cell populations in the zebrafish telencephalon as it undergoes the morphological changes underlying eversion. These datasets were registered to reference brains to form an atlas of telencephalic development at key stages of the eversion process (1dpf, 2dpf and 5dpf) and compared to expression in adulthood. Finally, we registered gata2:EGFP^bi105^ expression to the Zebrafish Brain Browser 6dpf reference brain (ZBB, see Marquart et al., 2015, 2017; Tabor et al. 2019), to allow comparison of this expression pattern with anatomical data already in ZBB.

## 2. Introduction

The telencephalon is the alar portion of the secondary prosencephalon (see Puelles and Rubenstein, 2003, 2015), consisting of the olfactory bulbs and the telencephalic lobes. It is responsible for higher brain functions, such as memory, emotions, cognition and higher-level multisensory integration and motor control. In zebrafish, as in all other ray-finned fish, the telencephalon is everted, a markedly different morphology to that of other vertebrates that have an evaginated telencephalon (Wullimann and Mueller, 2004b; Folgueira et al., 2012).

The morphological differences between everted and evaginated telencephali complicate comparisons and establishing homologies. In ray-finned fish, the telencephalon consists of two solid lobes separated by a midline ventricle that also extends over the dorsal surface under the *tela choroidea* (Butler, 2000; Yamamoto et al., 2007; Braford, 2009; Mueller and Wullimann, 2009; Nieuwenhuys, 2009a; Folgueira et al., 2012, Porter and Mueller, 2020). In other vertebrates, the telencephalon is evaginated and consists of two hollow hemispheres that surround an inflated, lateral ventricle. There is consensus that the dorsal and ventral portions of the everted telencephalon are homologous to the pallium and subpallium respectively, but establishing further homologies has proven to be challenging, especially for the pallium (Wullimann and Mueller, 2004b; Northcutt, 2008, Nieuwenhuys, 2009a; Mueller et al., 2011; Ganz et al., 2012; Porter and Mueller, 2020). This is because eversion is more complex than just a simple lateral out-folding of the neural tube (Yoshimoto et al., 2007; Mueller et al., 2011; Folgueira et al., 2012; Porter and Mueller, 2020), as first proposed by Gage (1893) and Studnicka (1894, 1896). For instance, based on the analysis of early forebrain development, we showed that there are two early morphogenetic events critical to formation of an everted telencephalon in zebrafish (Folgueira et al., 2012). Our model showed that eversion is more complex than a simple out-folding of the neural tube, but its implications are yet to be determined by detailed analysis of the developmental changes in locations of discrete cell populations during eversion (Briscoe and Ragsdale, 2019). After the early developmental events that generate an everted telencephalon, other processes, such as cell differentiation and migration, will certainly contribute to generate the organization of pallial subdivisions observed in the adult (Yoshimoto et al., 2007; Mueller et al., 2011; Porter and Mueller, 2020).

The principle aim of this study was to better understand how the organisation of the telencephalon changes during early development, when key events of eversion happen. To achieve this, we characterised the Et(*gata2:EGFP*)^bi105^ transgenic line (Folgueira et al., 2012; Turner et al., 2016), mapped the insertion site to *egr3* locus, examined *egr3* gene expression, and characterised the neurochemistry, cell cycle state and cell morphology of labelled EGFP positive (+) telencephalic cells. In combination with other well-characterized transgenic lines and structural markers, we tracked the distribution of various cell populations during early development. By combinatorial use of these diverse approaches we have generated a structural atlas of the developing zebrafish telencephalon, which should act as a framework, or scaffold to expedite further anatomical and functional studies.

## 3. Materials and Methods

### 3.1. Fish Stocks and Maintenance

Adult zebrafish (*Danio rerio*, Cyprinidae) were maintained under standard conditions that meet FELASA guidelines (Aleström et al., 2020): 28°C and 14 h light/10 h dark periods (Westerfield, 2000) at University College London (UCL) Fish Facility. Embryos were raised in fish water at 28°C and staged according to Kimmel et al. (1995). Phenylthiocarbamide (PTU, Sigma) at a concentration of 0.003% w/v was added to the fish water at 24 h post fertilization (hpf) to prevent pigment formation in larvae. All experimental procedures were conducted under licence from the UK Home Office, following UK Home Office regulations and/or European Community Guidelines on animal care and experimentation and were approved by animal care and use committees.

The following zebrafish strains were used in this study: AB and TL wild types, Et(*gata2*:*EGFP*)^bi105^ (Folgueira et al., 2012), Tg(*1.4dlx5a-dlx6a*:GFP)^ot1^ (Zerucha et al., 2000), Tg(*isl1*:GFP)^rw0^ (Higashijima et al., 2000) and Tg(-*10lhx2a:EGFP*)^zf176^ (Miyasaka et al., 2009; 2014).

### 3.2. BrdU Staining

BrdU (5-bromo-2’-deoxyuridine) incorporation assay was performed on 4dpf larvae by first embedding fish in 1% low-melting-point agarose and then injecting approximately 1nl of 10 mM BrdU (Sigma) into the heart. Larvae were then removed from the agarose and allowed to freely swim for 2 hours at 28.5°C. After anaesthesia and fixation, larvae were processed for immunohistochemistry (see below).

### 3.3 Anaesthesia and fixation

Specimens were deeply anesthetized in 0.2% tricaine methanesulfonate (MS222, Sigma) in fresh water and fixed in 4% paraformaldehyde (PFA) in phosphate buffered saline (PBS) with 4% sucrose by immersion. The tissue was then post-fixed in the same fixative for 24 h at room temperature. After this, cranial skin, jaw and other tissues were removed from embryos and larvae by fine dissection using two pairs of sharp forceps on embryos pinned in Sylgard (Turner et al., 2014). After washing in PBS and dehydration in methanol, they were kept at −20°C for at least 24 h. Brains from 20dpf fish and adults were dissected and kept in PBS at 4°C until use.

### 3.4. Immunostaining

Embryos and larvae were stained as whole mounts following standard procedures (Shanmugalingam et al., 2000; Turner et al., 2014). In brief, specimens were rehydrated to phosphate buffered saline with 0.5% Triton X-100 (PBS-T). After a brief proteinase digestion to improve permeability of the tissue, specimens were incubated with primary antibody overnight at 4°C. Specimens were then washed in PBS-T and incubated with fluorescent secondary antibodies again overnight at 4°C. After washing in PBS-T, embryos and larvae were mounted in 2% low-melting point agarose and imaged on a Leica confocal laser scanning system. For a detailed protocol on larval dissection, antibody labelling and mounting for imaging see Turner et al. (2014).

Brains from 20dpf fish were cryoprotected, frozen with liquid nitrogen-cooled methylbutane and cut in sections (12 µm thick) on a cryostat. After mounting the sections on gelatinized slides, they were rinsed in PBS-T and preincubated with normal goat serum (Sigma, 1:10) for 1 h. Next, they were incubated with a primary antibody overnight at room temperature and then with secondary fluorescent antibodies for 1 h, followed by PBS washes. Sections then stained for 5 minutes in a solution of Sytox Orange diluted in PBS-T, then mounted using glycerol-based mounting medium and photographed under a conventional epifluorescence microscope (Nikon Eclipse 90i).

For staining brains from 20dpf fish as whole mounts we followed the same protocol as for larval whole mount immunostaining, but with longer proteinase digestion and primary antibody incubation (48 h).

Antibodies and dilutions used were as follows:

#### Primary antibodies

Rabbit anti-green fluorescent protein (GFP; Torrey Pines Biolabs, Cat# TP401, dilution 1:1000), rat anti-GFP (Nacalai Tesque; Cat# GF090R, dilution 1:1000), mouse anti-bromodeoxyuridine (BrdU, Roche; Cat# B8434, dilution 1:300), mouse anti zona occludens 1 (ZO1, Invitrogen; Cat**#** 33-9100, dilution 1:300), mouse anti-acetylated tubulin antibody (IgG2b; α-tubulin; Sigma; Cat# T7451, dilution 1:250), rabbit anti-γ-aminobutyric acid (GABA; Sigma; Cat# A2052, dilution 1:1000), mouse anti-synaptic vesicle protein 2 (IgG1; SV2; DSHB; Cat# AB 2315387, dilution 1:250), rabbit anti-DSRed antibody (Living Colours; Clontech; Cat# 632496, dilution 1:300), rabbit anti-red fluorescent protein (RFP, MBL; Cat#PM005; dilution 1:2000).

#### Secondary antibodies

Alexa Fluor 488 (Invitrogen, Goat anti-Rabbit, Cat# A-11034, Goat anti-Rat Cat# A-11006, Goat anti-Mouse, Cat# A-11029, dilution all 1:200), Alexa Fluor 568 (Invitrogen, Goat anti-Mouse IgG, Cat# A-11031, Goat anti-Mouse IgG2b, Cat# A-21144, dilution all 1:200) and Alexa Fluor 633 (Invitrogen, Goat anti-Mouse IgG, Cat# A-21052; Goat anti-Mouse IgG1, Cat# A-21126, dilution all 1:200). To detect anti-acetylated tubulin and anti-SV2 in the same sample, isotype-specific secondary antibodies were used: Alexa Fluor 568 (IgG2b) and Alexa Fluor 633 (IgG1).

### 3.5. Whole mount fluorescent in situ hybridisation (FISH)

The FISH protocol was adapted from Jülich et al. (2005) and performed as described in Turner et al. (2016). Digoxigenin probes were made by standard protocols and were detected using the anti-DIG POD antibody (Roche, 1:1000) and stained using Cy3-tyramide substrate (Perkin Elmer, 1:50 in amplification buffer). After staining, antibody labelling for GFP was performed as above using rabbit anti-green fluorescent protein (GFP, Torrey Pines Biolabs, 1:1000) and goat anti-rabbit Alexa Fluor 488 (Invitrogen, 1:200) antibodies without further PK digestion. Cell nuclei were labelled with TOTO3-iodide (Invitrogen, 1:5000). Embryos were mounted in 1% agarose in 80% glycerol/PBS solution and imaged on a Leica SP8 confocal microscope.

### 3.6. Fluorescent Nucleic Acid Staining

Larval brains were whole mount stained with Sytox Orange (S11368; ThermoFisher) and used for studying cell architecture and distribution of different nuclei. Following antibody labelling, larval brains were washed in PBS and then stained for 30 minutes in a solution of Sytox Orange diluted in PBS-T (1:20,000). Subsequently, brains were washed in PBS and mounted for imaging.

### 3.7. Genome Walker mapping of enhancer trap insertion

The genomic location of the insertion in the enhancer-trap line Et*(gata2:EGFP)*^bi105^ was mapped using a linker-mediated approach with the Universal Genome Walker™ 2.0 Kit (Clontech). A detailed user manual is available online. Since the Et(*gata2*:*EGFP*)^bi105^ line was made using the Tol2-transposase system for genomic insertion (Kawakami et al., 2000, 2004). Using this information, two nested primers specific to the *tol2* sequence were designed: one for primary PCR (GSP1) and one for the secondary nested PCR (GSP2). These primers were non-overlapping, annealing to sequences as close to the end of the known sequence as possible. A combination of these primers (sequences below) and the ligated linkers/primer from the kit were used to generate four Genome Walker PCR amplicons from two restriction-generated genomic libraries. These amplicons were purified using Qiagen QIAquick PCR purification kit and checked for purity and concentration using a NanoDrop 2000C (ThermoScientific). Sanger-sequencing (Source BioScience, UK) using either stock primers such as M13F (within linker annealed kit constructs) or gene specific primers (see supplementary information for identified sequences) was checked for novel genomic sequence adjacent to the *tol2* sequence in each fragment, these sequences were aligned by BLAST/BLAT against the zebrafish genome. All four amplicons aligned to the same region of chromosome 8 (Fig. 1A). The insertion site was confirmed using PCR with primers specific to the specific EGFP employed in the transgenesis construct and the newly sequenced adjacent genomic sequence.

**Figure 1:**
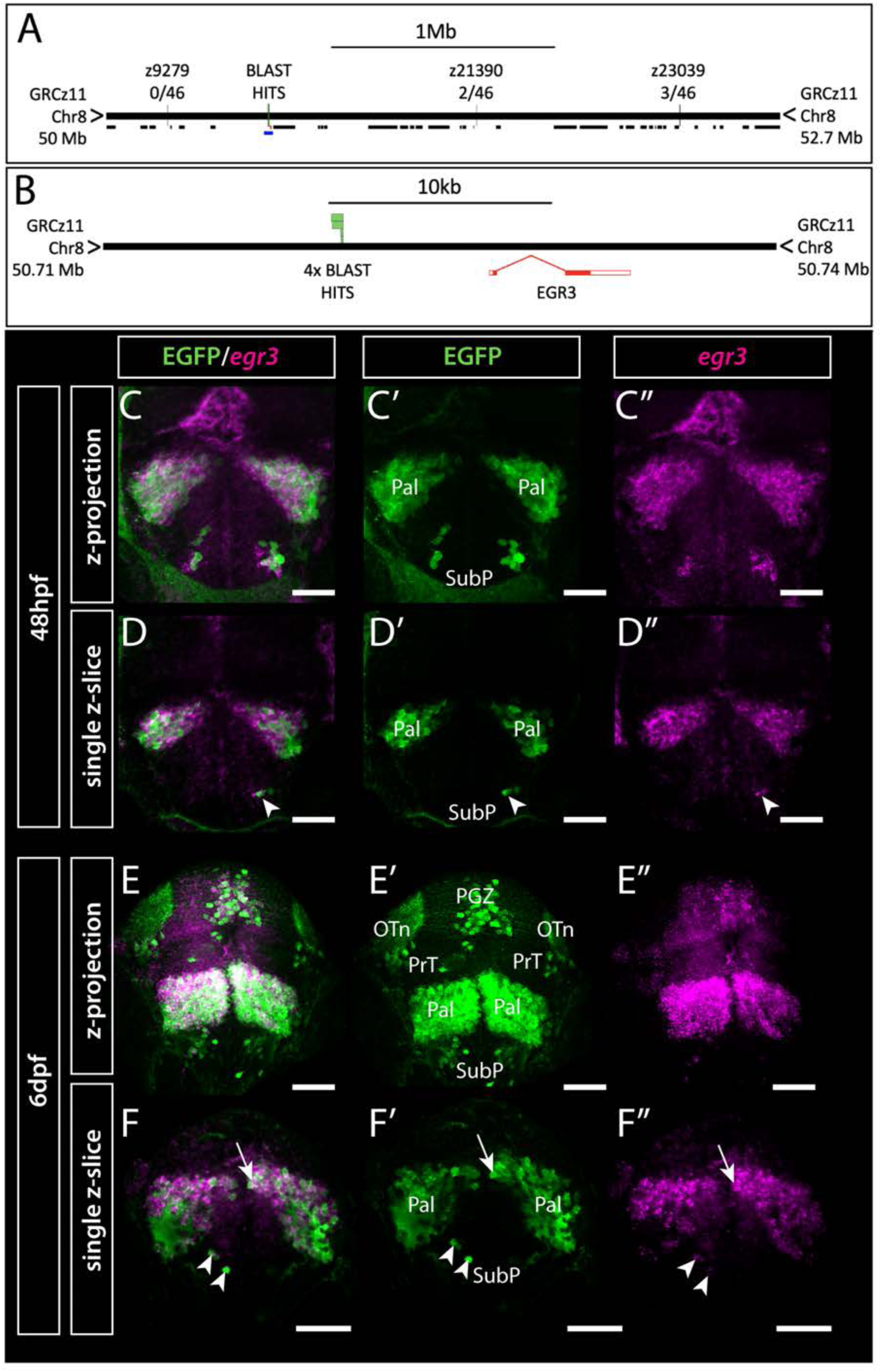
Et(*gata2:EGFP*)^bi105^ insertion is located near *egr3* and expression of *egr3* matches transgene expression. Mapping of Et*(gata2:EGFP)^bi105^* insertion showing the position of Genome Walker BLAST hits on assembly GRCz11, relative position of neighbouring genes (black lines) and linkage markers with meiotic recombination data **(A),** the lower blue line in **A** indicates the region around *egr3* shown in **B**, with BLAST hits (insertion) locations. **C-F”:** Immunostaining and *in situ* hybridization showing EGFP and *egr3* mRNA co-expression in the telencephalon and optic tectum of Et*(gata2:EGFP)^bi105^* fish at 48hpf **(C-D”)** and 6dpf **(E-F”)** in frontal view. C-C” and E-E” show z-projections and D-D” and F-F” show single z-slices. Note the co-expression of EGFP and *egr3* in pallial (arrows) and subpallial (arrowheads) cells. Scale bars: 50μm.

Nested sets of primers used with the adapter linked primers.

TTAAATTAAACTGGGCATCAGC, tol2_3 out_1
GGTTTGGTAATAGCAAGGGAAA, tol2_3 out_2
AAATACAAACAGTTCTAAAGCAGGA, tol2_5 out_1
CCTTGTATGCATTTCATTTAATGTT, tol2_5 out_2

### 3.8 Genetic linkage analysis

To confirm linkage between transgene and genome hits derived from Genome Walker use we employed traditional linkage mapping using simple sequence length polymorphisms (SSLPs)(Knapik et al., 1998). Briefly, nearby SSLPs were identified from zebrafish information network (ZFIN; zfin.org) webpages and Ensembl. A small panel of 48 transgene-positive, and 48 transgene-negative embryos were arrayed on a 96 well plate and genomic DNA was extracted by proteinase K digestion. PCRs for SSLPs were carried out on this arrayed DNA followed by gel electrophoresis. Where nearby SSLP markers were polymorphic between positive and negative groups, linkage was tested by measuring differential meiotic recombination between transgene positive and negative groups. The SSLP markers and primer sequences employed are listed below.

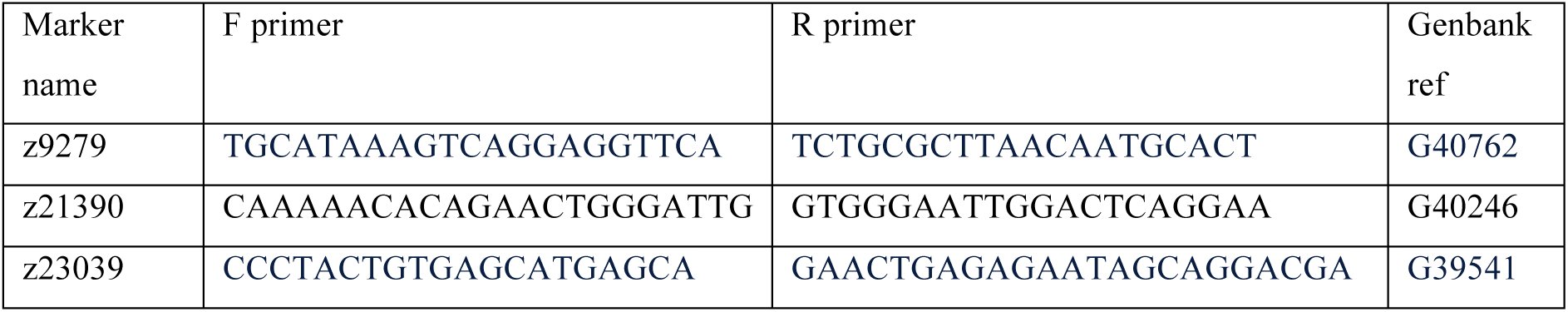

### 3.9. Preparation of sgRNAs, Cas9 mRNA

Template DNA for short guide RNAs (sgRNAs) synthesis was digested with DraI, and sgRNAs were transcribed using the MAXIscript T7 kit (Life Technologies). The pT3TS-nCas9n plasmid (Addgene) (Jao et al., 2013) was linearized with XbaI (Promega) and mRNA synthesised with the mMessage mMachine T3 Transcription Kit (Ambion). Transcription reactions were incubated in a water bath at 37°C for 2h or longer. To digest template DNA, 1 µL of TURBO DNase was added and reaction incubated for a further 15min at 37°C. The Cas9 mRNA was polyadenylated using the polyA tailing kit (Ambion). Cas9 mRNA and sgRNAs were purified using either the RNeasy Minikit (Qiagen) or Zymo columns.

### 3.10. Microinjections

Microinjection of DNA and RNA was performed using a borosilicate glass capillary needle attached to a Picospritzer III injector. Adult zebrafish were paired with dividers the night before injections. Dividers were removed the following morning and embryos were collected soon after to ensure they were at early 1 cell stage at time of injections. Embryos were aligned against a glass slide inside an upturned petri dish lid. The needle was calibrated to inject 1nl per embryo. Injections were performed into the cell for DNA and RNA.

### 3.11. Single cell labelling using EGFP to Gal4 switching with CRISPR/Cas9.

To achieve somatic switching of transgenes and mosaically label single or small groups of neurons within the Et(*gata2*:*EGFP*)^bi105^ transgenic line we adapted a technique developed by Auer et al. (2014) using the modified donor plasmid hs:Gbait (Kimura et al., 2014). This CRISPR/Cas9-mediated knock-in of DNA cassettes into the zebrafish genome uses homology-independent double-strand break repair pathways. Co-injection of a donor plasmid with a sgRNA and Cas9 nuclease mRNA results in concurrent cleavage of the donor plasmid and the selected chromosomal integration site resulting in the targeted integration of donor DNA. Using an EGFP target sequence in the donor plasmid and a sgRNA against EGFP Auer et al (2014) effectively converted EGFP transgenic lines into Gal4 versions. Kimura et al (2014) modified the donor plasmid (hs:Gbait) to include a heat-shock promoter to increase the efficiency of the switching of EGFP to Gal4, as the Gal4 cassette would be expressed regardless of which orientation the hs:GBait integrates into the genome.

To achieve sparse labelling of single EGFP expressing neurons in Et(*gata2*:*EGFP*)^bi105^ injected embryos we added 5xUASTdTom DNA to the injection mixture. This led to very mosaic labelling with TdTom of neurons within the original EGFP expression pattern. All plasmids, including hs:GBait (Kimura et al., 2014), were a kind gift from Thomas Auer. For the acute substitution, each embryo was injected with a solution containing 60pg of sgRNA EGFP1, 150pg of Cas9 mRNA, 7pg of Gbait-hsp70:Gal4 and 5pg UAS:TdTom (Zhang et al., 2012). Embryos were screened for RFP expression from 24hpf onwards. Specimens with desired expression were fixed and processed for immunohistochemistry. Technical limitations of this labelling technique, due to genome editing occurring early in development, made it challenging to label later born neurons.

### 3.12. Imaging

Fish were mounted in 2% low melting point agarose, either in lateral or dorsal view, under a dissecting scope equipped with bright field and fluorescent filters. Imaging was performed using either Leica SP2 or SP8 Confocal Microscopes equipped with filter-free AOBS systems tuned to the appropriate GFP/RFP spectra, with either water 25x (NA: 0.95) or 40x (NA: 0.8) immersion lenses and 3x line averaging. Imaging parameters varied for different data sets and scale references are included on figures to account for these differences. Due to the opacity of the tissue, imaging beyond a total depth of 200-300 µm from the surface of the fish brain was not possible.

### 3.13. Image analysis and Editing

Confocal stacks were processed using Fiji (ImageJ) and/or Volocity and/or Imaris. Images and figures were assembled using Adobe Photoshop or Adobe Illustrator. Unless otherwise stated, the nomenclature used in this study is largely based on that of the adult zebrafish brain atlas (Wullimann et al., 1996) and early zebrafish brain development (Mueller and Wullimann, 2005). For larval optic tectum nomenclature, we followed Robles et al. (2011).

### 3.14. Image registration

Registration of image volumes was performed using the ANTs toolbox version 2.1.0 (Avants et al., 2011) using UCL’s computer cluster, typically with a Dell C6220 node with 16 cores and 16 GB of RAM. Individual brain volumes for each age and orientation (lateral or dorsal view) were registered using as template either the ZO1, acetylated tubulin, or both imaging channels of an example fish in each category. All registrations were manually assessed for global and local alignment accuracy.

As an example, to register the 3D image volume in fish1–01.nii to the reference brain ref.nii, the following parameters were used: antsRegistration -d 3 –float 1 -o [fish1_, fish1_Warped.nii.gz] –n BSpline -r [ref.nii, fish1– 01.nii,1] -t Rigid[0.1] -m GC[ref.nii, fish1–01.nii,1,32, Regular,0.25] -c [200x200x200x0,1e-8,10] –f 12x8x4x2 –s 4x3x2x1-t Affine[0.1] -m GC[ref.nii, fish1–01.nii,1,32, Regular,0.25] - c [200x200x200x0,1e-8,10] –f 12x8x4x2 –s 4x3x2x1-t SyN[0.1,6,0] -m CC[ref.nii, fish1– 01.nii,1,2] -c [200x200x200x200x10,1e-7,10] –f 12x8x4x2x1 –s 4x3x2x1x0

The deformation matrices computed above were then applied to any other image channel N of fish1 using:

antsApplyTransforms -d 3 -v 0 –float -n BSpline -i fish1–0N.nii -r ref.nii -o fish1– 0N_Warped.nii.gz -t fish1_1Warp.nii.gz -t fish1_0GenericAffine.mat

A registered 6dpf Et(*gata2*:*EGFP*)^bi105^ stack (.PNG file accompanying this article, see supplementary data) can be viewed in the online Zebrafish Brain Browser browser (Marquart et al., 2015; Marquart et al., 2017) at http://metagrid2.sv.vt.edu/~chris526/zbb/. To add any ZBB registered stack, go to “Lines” and select “Custom”. From there you can “Select File” to select the .PNG file. Click “Load” and the stack should now display in the browser, which you can then visualise with any other ZBB registered line or label.

### 3.15 Terminology

According to the neuromeric model (Puelles and Rubenstein, 2003, 2015), the neuraxis is bent so the optic chiasm is the anterior tip of the brain. Based on this model, the alar telencephalon would be divided in an anterior subpallium and a posterior pallium (see Herget et al., 2014). The telencephalon, however, has been classically divided in a dorsal division or pallium and a ventral division or subpallium. Thus, for clarity in our anatomical descriptions, we use body axis coordinates (“rostral”-towards the nose; “caudal”-towards the tail; “dorsal” and “ventral”) for brain structures (Fig. 2, top schematic) (see also Herget et al., 2014 and Porter and Mueller, 2020). We use the terms “alar” and “basal” whenever appropriate (Herget et al., 2014; Porter and Mueller, 2020) (Fig. 2, top schematic).

**Figure 2:**
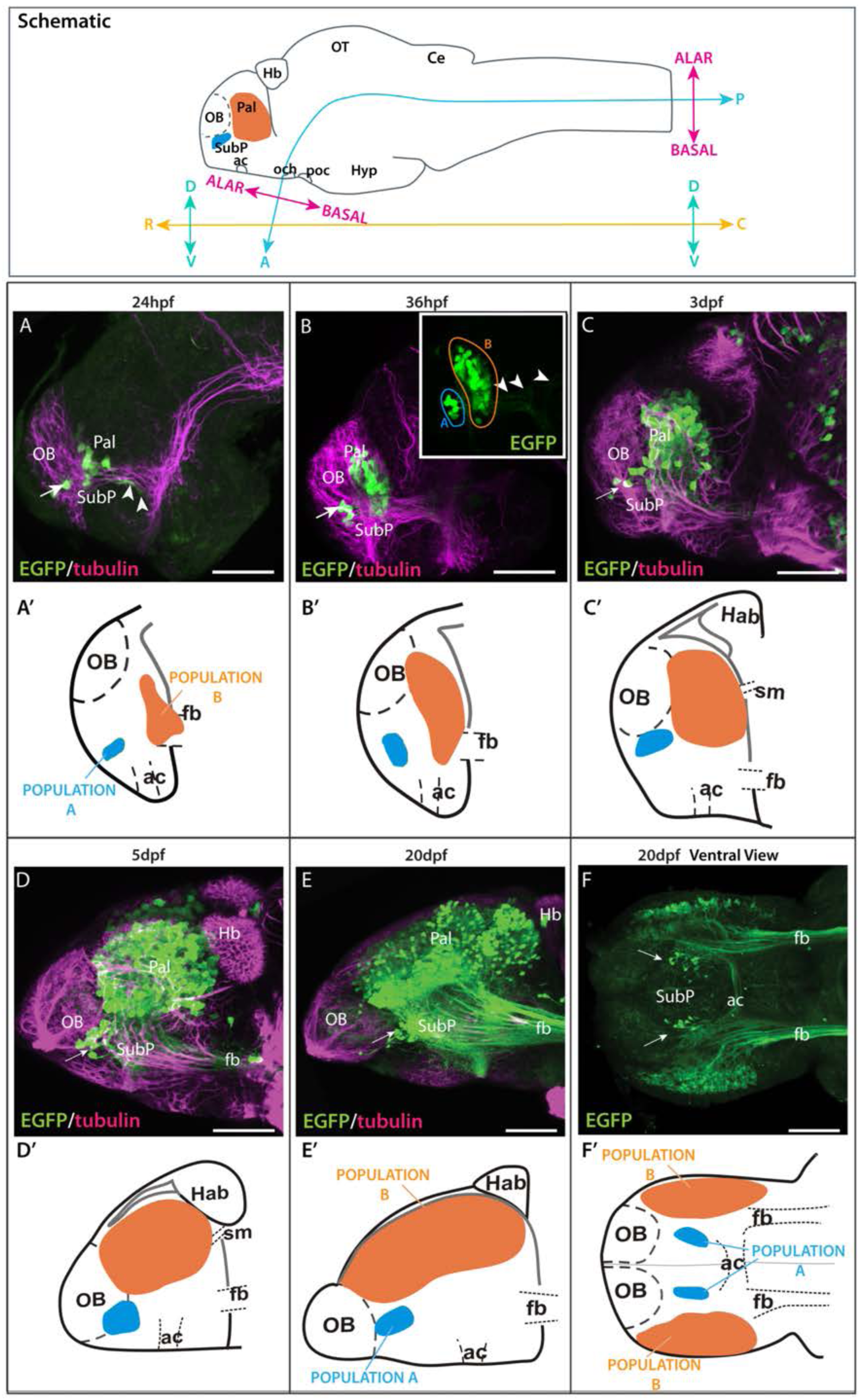
Time series showing Et(*gata2:EGFP*)^bi105^ expression. Top box: sagittal view of a 3dpf zebrafish larval brain illustrating the body axis coordinates (“rostral”, “caudal” “dorsal” and “ventral”) predominantly used in this paper (adapted from Herget et al., 2014). For reference, the bent neuraxis of the neural tube is also shown (see Herget et al., 2014) with the Alar/Basal subdivision of the brain defined according to the neuromeric model (Puelles and Rubenstein, 2003, 2015). Lateral **(A-E)** and ventral **(F)** views of zebrafish brains stained against EGFP only (**B inset**, **F**) or EGFP in combination with acetylated tubulin **(A-E)** from 24 hpf to 20dpf. Arrow points to the ventral EGFP+ population identified as a subpallial nucleus. Arrowheads in **A** and **B inset** point to fibres exiting the telencephalon along the forebrain bundle towards caudal areas. **A’-F’**: Schematics showing the changes in location of the EGFP+ populations (A and B) relative to telencephalic regions and tracts over time. Rostral to the left. Scale bars: **(A-D)** 50μm; **(E,F)** Abbreviations: A, anterior; C, caudal; D, dorsal; P, posterior; V, ventral. 100μm.

## 4. Results

### 4.1. The *Et*(*gata2*:*EGFP*)^bi105^ transgene neighbours *early growth response 3 (egr3)* and EGFP expression closely matches endogenous *egr3* expression

EGFP is expressed in the telencephalon of Et(*gata2*:*EGFP*)^bi105^ larvae and is a useful marker to understand telencephalic morphogenesis (Folgueira et al., 2012; Turner et al., 2016). The genomic location of this transgene was, however, not known.

To identify this locus, we mapped the position of the Et(*gata2*:*EGFP)*^bi105^ insertion using linker-mediated PCR, confirmed by genetic linkage mapping. Our results showed that the genomic insertion of the transgene in Et(*gata2*:*EGFP*)^bi105^ fish lies 6,543bp upstream of the first exon of *Early Growth Response 3* (*egr3)* (see methods and Fig. 1A-B) which encodes a transcription factor that functions as an immediate-early growth response gene in some systems (Yamagata et al., 1994).

With the insertion point established, we then assessed whether Et(*gata2*:*EGFP*)^bi105^ transgene expression recapitulates *egr3* expression. We examined *egr3* mRNA expression using fluorescent *in-situ* hybridisation in combination with immunohistochemistry for EGFP in Et(*gata2*:*EGFP*)^bi105^ embryos. There was extensive co-localisation of EGFP protein and *egr3* mRNA in the telencephalon from 48hpf (Fig. 1C-D’’). At this stage, there is also strong *egr3* mRNA expression in the midline of the optic tectum, but little EGFP protein observed in this location (Fig. 1C-C”, D-D’’). By 6dpf, *egr3* mRNA expression overlaps with EGFP protein in the pallium, subpallium and optic tectum of Et(*gata2*:*EGFP*)^bi105^ embryos (Fig. 1E-F’’).

In summary, this demonstrates that the Et(*gata2*:*EGFP*)^bi105^ transgene insertion maps near *egr3,* and EGFP expression is nested within the endogenous *egr3* expression.

### 4.2. Changes in telencephalic topography between 18hpf and 20dpf

We used wholemount immunohistochemistry against EGFP in Et(*gata2*:*EGFP*)^bi105^ fish from 18hpf to 20dpf to follow the development of pallial and subpallial telencephalic populations, combined with anti-acetylated tubulin to highlight the progressive development of major axon pathways (Fig. 2A-F’; Wilson et al. 1990; Chitnis and Kuwada 1990). Following EGFP expression over time does not allow us to definitively identify the same cells at each stage, but our observations strongly support there being stable transgene expression within the same discrete telencephalic cell populations over time.

By 20hpf, the first few EGFP+ cells were evident ventrally in the telencephalon (not shown), dorsal to the anterior commissure and forebrain bundle (telencephalic tract of Chitnis and Kuwada, 1990; supraoptic tract of Wilson et al., 1990). By 24 hpf, EGFP+ cells are divided in two populations (Fig. 2A): “population A” is located just rostral to the anterior commissure (subpallial nucleus, see below) and “population B” is located more caudally and dorsally (pallial population, see below). At this stage, EGFP+ axons and growth cones from these neurons were observed within the forebrain bundle (Fig. 2A).

By 36hpf, an increase in EGFP+ cells was evident in both populations (Fig. 2B), especially in “population B” (pallial population). The cells of “population A” (subpallial population) extended processes to the dorsal part of the anterior commissure (Fig. 2B-C), where they intermingled with processes from the EGFP+ cells of “population B”. Some fibres extended along the forebrain bundle towards caudal forebrain areas (Fig. 2B inset).

To confirm the regional identity of the EGFP+ telencephalic cells, Et*(gata2:EGFP)*^bi105^ embryos were labelled with immunohistochemisty against EGFP and fluorescent *in situ* hybridisation against *dlx1a* and *tbr1* at 36 hpf (MacDonald et al., 2010) (Fig. 3). *tbr1* encodes a transcription factor that marks developing glutamatergic cells (Hevner et al., 2001, 2006; Puelles et al., 2000; Englund et al., 2005). In zebrafish, *tbr1* is mainly expressed in the pallium, with two smaller additional expression sites in the septal areas and in telencephalic populations that derive from the prethalamic eminence (Mione et al., 2001; Wullimann and Mueller., 2004a,b; Mueller et al., 2008, Turner et al., 2016). *dlx1a* is expressed in proliferative and immature neurons of the subpallium and preoptic area (MacDonald et al., 2010; Wullimann and Mueller, 2004a, b). Using these markers we found that for “population B”, EGFP+ cells were nested within the *tbr1+* domain (Fig 3A-A’’ and E-E’’) and were predominantly *dlx1a* negative (-) (Fig. 3B-B’ and F-F’). These observations confirmed the pallial character of this population. The transgene does not label the full set of pallial cells, as there are cells that are *tbr1+* but EGFP-, including the ventricular zone (Fig. 3A-A’’ and E-E’’). Single confocal slices through 36hpf larvae showed the EGFP+ cells of “population A” at the border of *tbr1+* domain (Fig. 3C-C’’ and G-G’’), but they are nested within the subpallial marker *dlx1a* (Fig. 3D-D’’ and H-H’’). The location of “population A” in relationship to *tbr1* and *dlx1a* domains confirms its subpallial identity.

**Figure 3:**
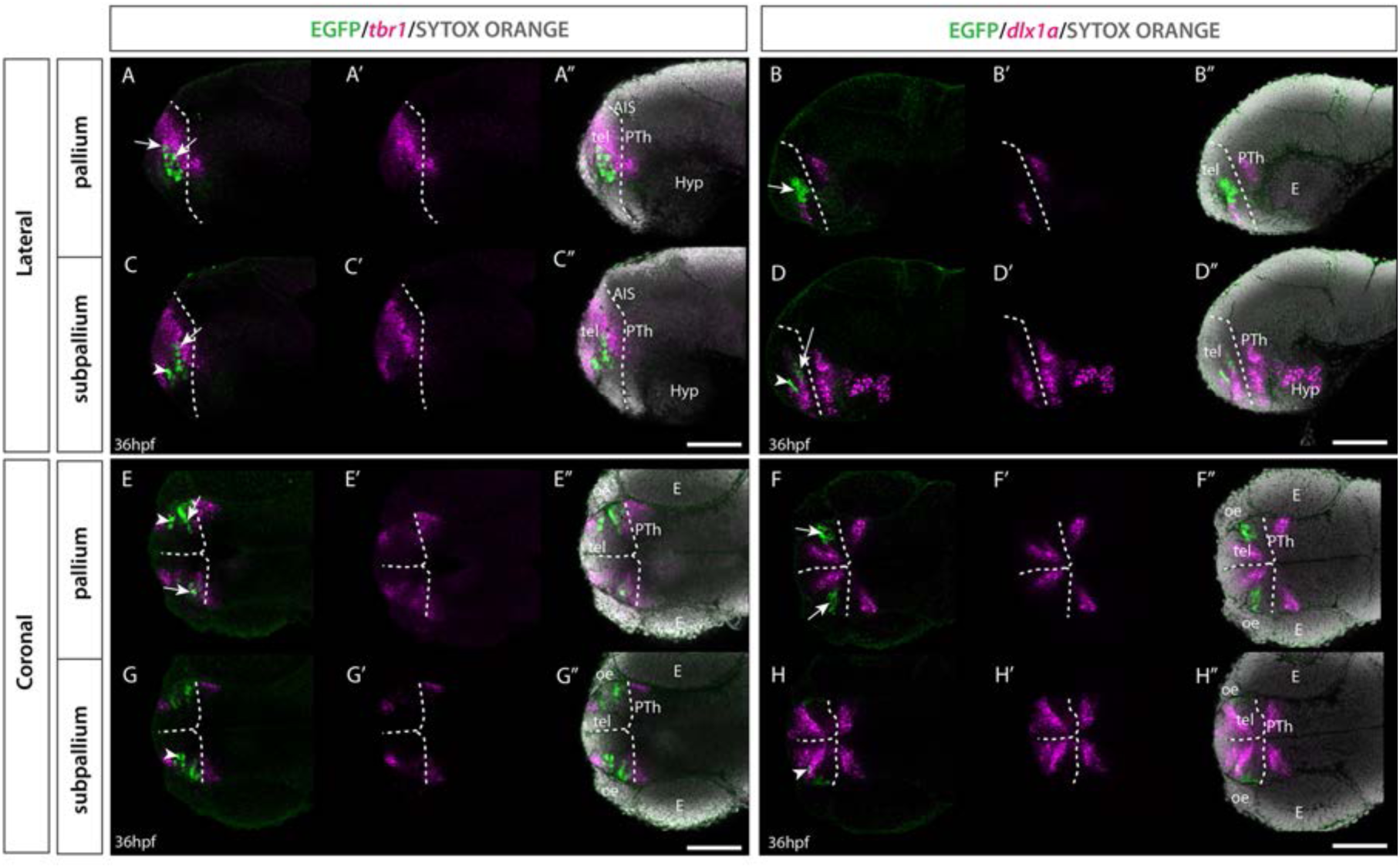
Et(*gata2:EGFP*)^bi105^ line expression in combination with *tbr1* and *dlx1a* markers. Sagittal (**A-D’’)** and horizontal **(E-H’’)** sections through a 36hpf Et*(gata2:EGFP)*^bi105^ fish labelled with anti-GFP (green) and FISH for *tbr1* (pink)**(A,A’’,C,C’’,E,E’’,G,G’’)** or *dlx1a* (pink) **(B,B’’,D,D’’,F,F’’,H,H’’)** with cell nuclei stained with sytox orange (grey). All images are single confocal z-slices taken from volumes, rostral to the left. Scale bars: 50μm.

From 36hpf to 3dpf (Fig. 2B-C) the number of EGFP+ cell bodies again increased in the pallial EGFP+ population. By 3dpf, a few EGFP+ fibres (but no cell bodies) were observed in the inner layer of the olfactory bulb. From 3dpf to 5dpf, the number of EGFP+ pallial cell bodies in the telencephalon increased substantially, as did the overall size of the pallial territory (Fig. 2C-D).

From 5dpf to 20dpf, we observed a dramatic increase in the size of the telencephalic pallial lobes relative to other neighbouring areas (Fig. 2E-F), such as the olfactory bulb and habenula. Although the EGFP+ subpallial cells were initially positioned rostral to the EGFP+ pallial cells, by 20dpf they sit ventral to this cell population (Fig. 2E-F). Large numbers of EGFP+ fibres were observed coursing towards caudal areas via the forebrain bundle (Fig. 2F) and crossing the midline in the anterior commissure (Fig. 2F). Fibres originating from the subpallial population were also observed in the anterior commissure at this stage (Fig. 2E-F).

The changes in the relative positions and sizes of the subpallial and pallial EGFP+ populations over time (from 24hpf to 20dpf) reflect the morphological changes that occur as a result of eversion (24hpf to 5dpf) and telencephalic growth (5dpf to 20dpf) (Folgueira et al., 2012; Dirian et al., 2014; Furlan et al., 2017)

As the anatomical divisions between regions are clearer by 20dpf, we could more precisely determine the identity of the different EGFP+ populations at this stage. Rostrally, many EGFP+ fibers (but no cell bodies) were present in the internal cell layer of the olfactory bulbs (Fig. 4A-B). In the telencephalic lobes, we observed that EGFP expression was not consistently strong throughout the dorsal telencephalon/pallium (Fig. 4B-E). Following nomenclature of Castro et al. (2006; see also Yañez et al., 2021), the principal areas of EGFP expression in the pallium were the medial zone (Dma and Dm of Castro et al., 2006), lateral zone (Dl) (Fig. 4B-E), central zone (Dc) (Fig. 4C-D) and scattered cells in the dorsal zone (Dd) (fig. 4C-E). In contrast to 4dpf larval expression (apparently in the whole pallium, excluding pallial derived olfactory bulb cells), we found very few EGFP+ cells in the anterior region of the lateral zone (Dla) and in the posterior zone (Dp) (Fig. 4B, E). EGFP expression was also excluded from the pallial ventricular zone (Fig. 4B-E). This suggests that EGFP expression is excluded from radial glia and proliferative cells which occupy these areas. A few EGFP+ cells were found in the lateral subpallium and close to the lateral forebrain bundle (lfb) (Fig. 4C, inset i). Based on this location, these subpallial cells could be part of the lateral nucleus of the ventral telencephalon (Vl; see Mueller et al., 2008; Mueller and Guo, 2009; Ganz et al., 2012).

**Figure 4:**
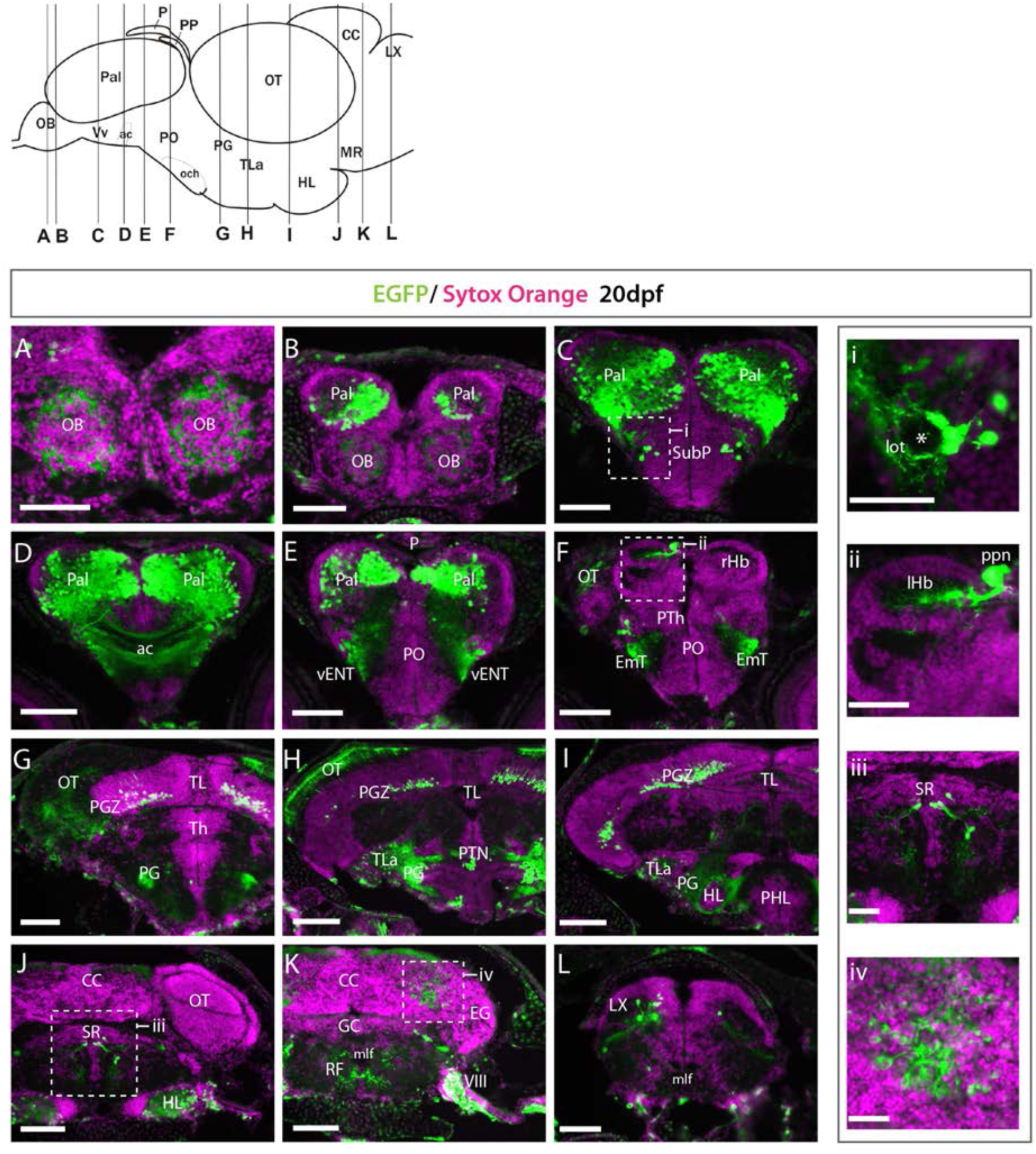
Et(*gata2:EGFP*)^bi105^ expression at 20dpf. A-L: Transverse section through an Et(*gata2*:EGFP)^bi105^ fish showing EGFP expression from rostral **(A)** to caudal areas **(L).** Schematic of lateral view of the brain (top left) shows levels of the sections. i-iv: detail of the areas marked in **C, F**, **J** and **K**. Areas of EGFP expression at this stage was consistent with those seen at 4dpf; the only *de novo* expression observed was in the cerebellum and vagal lobe **(K, L, iv)**. Scale bars: **(A-L)** 100μm; **(i-iv)** 40 μm.

### 4.3. Et*(gata2:EGFP^bi105^)* drives EGFP expression in post-mitotic glutamatergic telencephalic neurons

The position of EGFP+ cells distant from the ventricular zone (Fig. 5A-C), suggested that they are post-mitotic neurons. To ascertain if this is correct, we treated Et*(gata2:EGFP)*^bi105^ larvae with short pulses of BrdU at 4dpf and then performed immunostaining against EGFP and BrdU. The gata2:EGFP+ cells did not incorporate BrdU and, consequently, they are most probably non-proliferative, post-mitotic neurons (Fig. 5D-D’),

**Figure 5:**
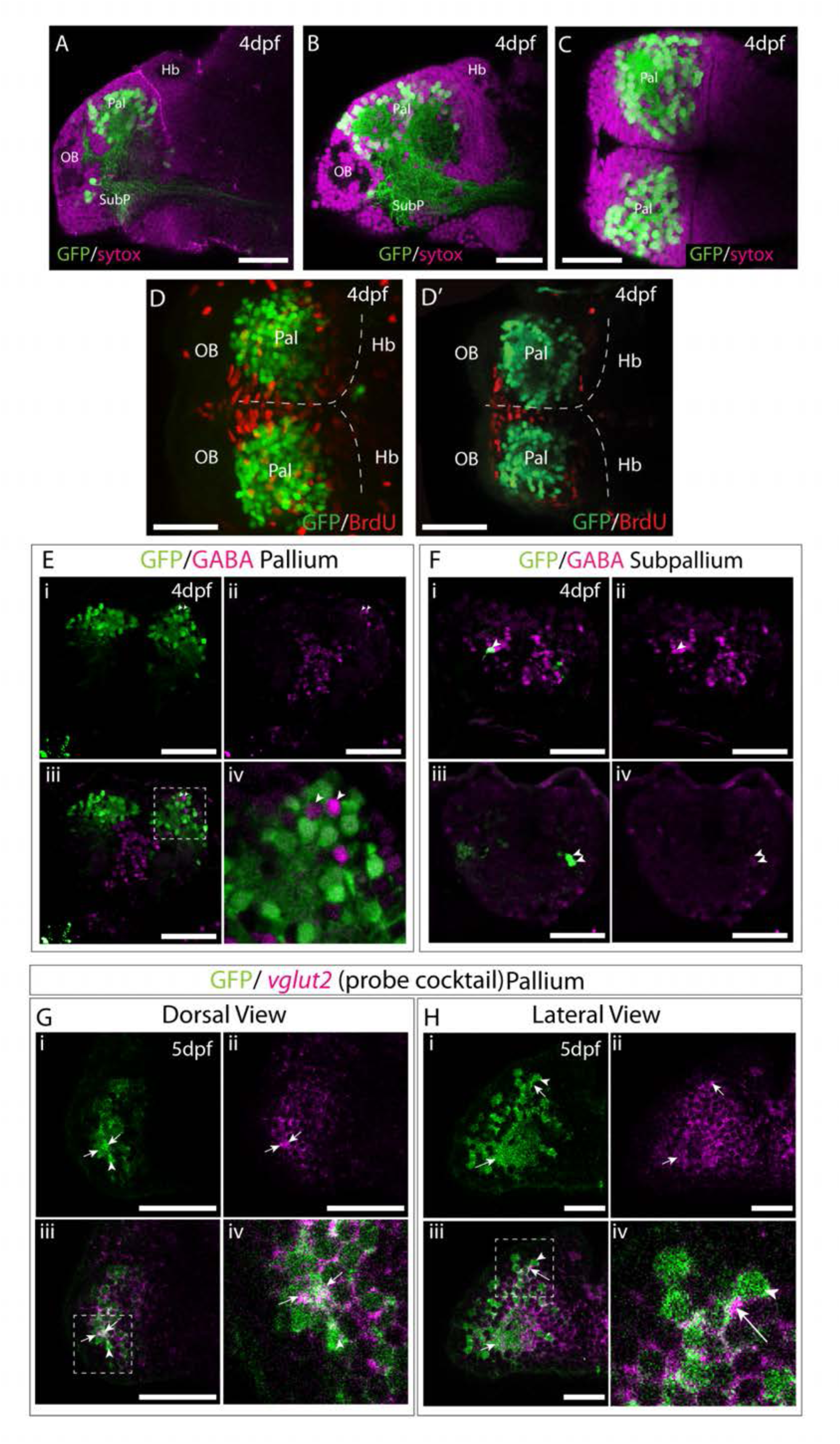
Et(*gata2:EGFP*)^bi105^ is expressed in post-mitotic glutamatergic telencephalic neurons. Sagittal **(A-B)** and horizontal **(C)** sections through a 4dpf Et*(gata2:EGFP)*^bi105^ brain stained with sytox (magenta) to label cell nuclei and anti-EGFP (green). EGFP expression is largely excluded from the dorsal and medial ventricular zones of the telencephalon. **D-D’:** Dorsal views of a 4dpf Et*(gata2:EGFP)* ^bi105^ brain labelled with anti-BrdU (red) and anti-GFP (green) antibodies. **(D)** z-projection and **(D’)** shows a single coronal z-slice through the same brain. BrdU is excluded from the EGFP+ neurons in the dorsal telencephalon indicating that these cells are likely to be non-proliferative and postmitotic. **E-F:** Transverse sections through the telencephalon of a 4dpf Et*(gata2:EGFP)* ^bi105^ brain stained with anti-GABA (magenta) to label GABAergic cells and anti-GFP (green). EGFP+ cells in the pallium **(E)** are for the most part GABA-. Gaps where GABAergic pallial interneurons intermingle with EGFP+ pallial neurons are visible (arrowheads in **Ei**). In subpallial telencephalic domains **(F)** some cells do co-express EGFP and GABA (arrowheads in **Fi-iv**). **Fi-ii** and **Fiii-iv** are different z-sections through the same Et*(gata2:EGFP)* ^bi105^ brain showing subpallial EGFP+ neurons on both sides of the brain. Dorsal **(G**) and lateral **(H)** views of Et*(gata2:EGFP)* ^bi105^ brains at 5dpf labelled with FISH using a “cocktail” probe for *vglut2a/slc17a6b* and *vglut2b/slc17a6a* (magenta) and antibody labelling with anti-GFP (green) show overlap of *vglut2a* and EGFP within the telencephalic neuropil (arrows) associated with EGFP+ cells (arrowheads). **Giv:** High-magnification view of dorsal telencephalic neurons in boxed region in **Giii**. **Hiv:** High-magnification view of dorsal telencephalic neurons in boxed region in **Hiii**. Scale bars: 50μm.

To further characterise the EGFP+ cells, we assessed if they are glutamatergic and/or GABAergic. Co-immunostaining against GFP and GABA showed no co-expression in the pallium at 4dpf (Fig. 5E). GABAergic cell bodies were intermingled with the EGFP+ cell bodies in a salt and pepper manner (Fig. 5iii-iv). Some EGFP+ cells in the subpallium were GABAergic (arrowheads in Fig. 5H). GABAergic neurons in the pallium are likely migrated interneurons (Wullimann and Mueller, 2004b; Mueller et al., 2006) and our results confirm that EGFP is not expressed in GABAergic pallial interneurons.

To test whether the EGFP+ cells are glutamatergic, a cocktail of probes against *vglut2a/slc17a6b* and *vglut2b/slc17a6a* (see Higashijima et al., 2004) was used for fluorescent *in situ* hybridisation on 4dpf Et*(gata2:EGFP)*^bi105^ embryos. This probe mix labels neurons in the forebrain very broadly, but areas of strong GABAergic expression, such as the subpallium, preoptic region and prethalamic areas lack staining. EGFP protein in the telencephalic neuropil (arrows) surrounding GFP+ cells (arrowheads) co-localises with *vglut2* mRNA (Fig. 5G-H). No or very faint *vglut2* mRNA expression was noted in subpallial cells (not shown).

In summary, these results show that EGFP+ telencephalic cells are post-mitotic, and based on cell morphology and location, are most likely to be differentiated neurons. The EGFP+ pallial population contain glutamatergic neurons, while the subpallial EGFP+ cluster contains GABAergic neurons.

### 4.4. Individual morphologies of EGFP+ neurons in Et*(gata2:EGFP)^bi105^* larvae

The density of pallial EGFP expression in Et*(gata2:EGFP)*^bi105^ larvae prevents any analysis of the morphology and projections of individual neurons. To visualise sparsely labelled neurons, we employed the CRISPR/Cas9 approach developed by Auer et al. (2014) and further modified by Kimura et al. (2014) to excise the EGFP transgene and mosaically insert Gal4. We modified this method slightly by adding a UAS:TdTomato construct to the injection mix. The mosaic inheritance of this DNA construct, layered upon the already mosaic conversion to Gal4, together with fine process staining properties of TdTomato, permitted both increased mosaicism and better labelling of neuronal processes (Schematic Fig. 6)

**Figure 6:**
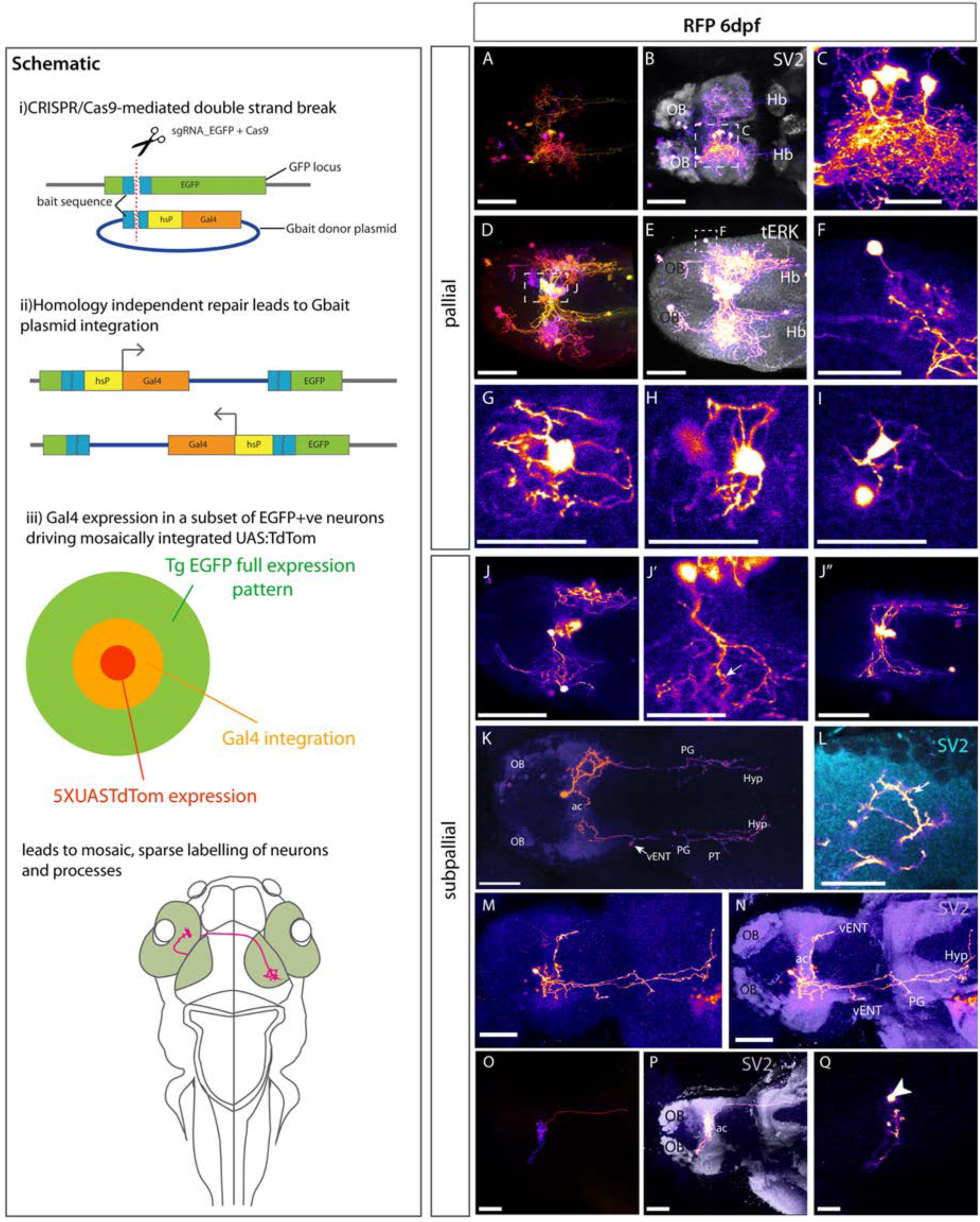
Morphologies of pallial and subpallial neurons. **Schematic:** Schematic parts **(i)** and **(ii)** adapted from Kimura et al., 2014 explaining the Crispr/Cas9 mediated method to switch EGFP to Gal4. **(iii)** Shows our adaptation of this method to sparsely label single neurons and their processes by adding 5XUASTdTom to the injection mix. **A-Q:** Et*(gata2:EGFP)*^bi105^ larvae injected with CRISPR cocktail and labelled with anti-RFP and other markers. **A-I: Pallial Neurons**. **(A-I)**. Dorsal views of 6dpf Et*(gata2:EGFP)* ^bi105^ larvae injected with CRISPR cocktail. Labelled with anti-RFP (FIRE depth colour code) **(A, D)** and FIRE LUT **(B-C, E-F)**, anti-SV2 (grey) **(B)**, anti-tERK (grey) **(E)**. **C:** Detail of “monopolar” dorsal pallial neurons from larva shown in **A**-**B** (area indicated by whisker box in **B**). **F:** Detail of single monopolar pallial neuron from larva shown in **D**-**E** (area indicated by whisker box in **E). G-I:** Three examples of basket shaped cells all found in the dorsal pallium of larva shown in **D-E**. **J-Q**: **Subpallial neurons**. **J-J”:** A cluster of midline subpallial cells from larva shown in **D-E** exhibiting elaborate dendritic trees that project throughout the subpallium at the level of the ac. These dendrites exhibit a spiny morphology (arrow in **J’**). **K-N:** Ventral view of two different 6dpf, labelled with anti-RFP (FIRE) and anti-SV2 (cyan in **L,** purple in **N**). **K:** Subpallial neuron with cell body just rostral to ac and bilateral projections. Arrow in **(K)** shows branching in the area of the lfb adjacent to the vENT. This cells also extends processes to tuberal and hypothalamic areas.. **L:** Close-up details of cell processes and dendritic spines (arrow) in the ipsilateral telencephalon from cell shown in **K.** **M-N:** Subpallial neuron also with cell body rostral to ac. It shows elaborated dendrites in the ipsilateral subpallium and one process crossing in the ac to innervate the contralateral telencephalon. Long processes project caudally to likely innervate the ventral entopeduncular nucleus, tuberal areas and hypothalamus. **O-Q:** A subpallial neuron with a peculiar cocoon morphology [ventral view; FIRE depth colour code in **O**, FIRE LUT in **P-Q,** anti-SV2 (grey) in P]. The cell body (arrowhead in **Q**) is located within the neuropil area of the ac. The dendrites wrap around the cell within the ac (**O**-**P**). A short process innervates the contralateral subpallium rostral to the ac (**P**). A longer process descends through the fb to contralateral tuberal regions (**P**). Scale bars **(A,B,D,E,J-J”,K,M,N,O,P,Q)**: 50μm. Scale bars **(C,F,G,H,I,L):** 25μm.

Injection of the CRISPR/Cas9/EGFP guideRNA/Gal4/UAS cocktail into one cell stage Et*(gata2:EGFP)*^bi105^ embryos lead to mosaic RFP labelling of small numbers of pallial neurons (Fig. 6A-I). All observations of morphology were made at 6 dpf, when the eversion process is almost complete. Labelled pallial neurons fell into two obviously distinct morphologies:

1. A population of neurons surrounding a central core of telencephalic neuropil (demarcated by SV2 expression, Fig. 6B). These cells had monopolar morphology with their single process extending towards the neuropil core before ramifying into multiple dendrites that formed a dense tangle with dendrites of adjacent neurons. Some labelled processes were also present in the contralateral telencephalic neuropil core (Fig. 6B). We were unable to ascertain whether these contralateral processes originated from these monopolar pallial neurons, perhaps crossing the anterior commissure, or other subpallial cells labelled in this specimen. Some processes also projected to caudal regions through the forebrain bundle. A survey of the single neuron morphologies registered to the mapzebrain atlas (Kunst et al., 2019) with soma located in the pallium (n=28) revealed some pallial neurons projecting through the forebrain bundle to innervate caudal regions, such as the hypothalamus. None of the pallial neurons in this mapzebrain dataset had processes that crossed in the anterior commissure to elaborate processes in the contralateral pallium. So, although this cannot be definitively ruled out, it seems more likely such processes originate from subpallial gata2:EGFP*+* neurons.
2. A population of multipolar neurons with stellate morphology (Fig. 6G-I). These neurons were positioned either at the dorsal surface or within the central neuropil core. These cells extended multiple processes directly from the cell body that appeared to encase small areas of neuropil or groups of cell bodies (Fig. 6G-I).

These experiments also allowed us to analyse the morphology of subpallial cells (Fig. 6J-Q). Figure 6K and L shows one such neuron whose cell body lay just rostral to the anterior commissure. This neuron was monopolar, with an elaborate dendritic arbour, some with spines, in the vicinity of the anterior commissure (Fig. 6K-L). The cell extended a bifurcated axon bilaterally down the left and right forebrain bundles to reach downstream targets that may include the ventral entopeduncular nucleus (vENT, arrow Fig. 6K), preglomerular, tuberal and hypothalamic areas (Fig. 6K). While the dendrites of this cell ramified only ipsilaterally, axon collaterals were bilateral and overtly symmetrical. Another example of a subpallial projection neuron with predominantly ipsilateral projections (Fig 6M-N) innervated similar caudal regions. Processes from this neuron also crossed the midline in the anterior commissure to terminate in the vicinity of the contralateral ventral entopeduncular nucleus. Axonal processes also crossed the midline in the hypothalamus. The subpallial neurons shown in Fig. 6J-J”, with cell bodies located at the midline, showed similar innervation of commissural regions and long descending axons. We observed one cell within the anterior commissure (Fig. 6O-Q). This cell exhibited cocoon-like dendrites, tightly coiling within the ac itself. It sent one process rostrally into the contralateral pallium at the border with the olfactory bulb and another caudally through the lateral forebrain bundle to the posterior tuberculum (PT).

A survey of single neuron morphologies registered to mapzebrain atlas (Kunst et al., 2019) with somata located in the subpallium (n=139) showed long-projecting neurons comparable to those shown in 6K-M to be very common with around 70% projecting either bilaterally or ipsilaterally to caudal regions, predominantly the hypothalamus, interpeduncular nucleus, posterior tuberculum and raphe nuclei. Unlike pallial cells in mapzebrain, the subpallial neurons frequently innervate the contralateral subpallium and pallium via the anterior commissure.

Mosaic analysis of neuron morphologies in Et*(gata2:EGFP)*^bi105^ larvae demonstrated that, even at larval stages, telencephalic cells already have elaborated intricate processes and dendritic arbors.

### 4.5. Registration of transgene expression reveals the morphogenetic rearrangements of telencephalic regions during development

As we have described, the Et*(gata2:EGFP)*^bi105^ transgene labels post-mitotic pallial cells, as well as a small group of subpallial cells. The restricted pallial expression of EGFP makes it a useful tool to track telencephalic cell populations through key stages of telencephalic development, particularly relating to eversion.

To build on our previous analyses of pallial eversion in zebrafish (Folgueira et al., 2012), we have taken advantage of non-linear volumetric image registration to register brains with labelling of different cell populations/structures to reference brains (Figures 7 and 8; Marquart et al., 2015, 2017; Randlett et al., 2015). We used acetylated tubulin labelling, common between all datasets, as a reference channel (Fig. 7E, K, Q; not shown in 8). These registered datasets describe how major telencephalic cell populations/structures (olfactory bulb, pallium, subpallium, and *tela choroidea*) change position relative to each other during development.

**Figure 7:**
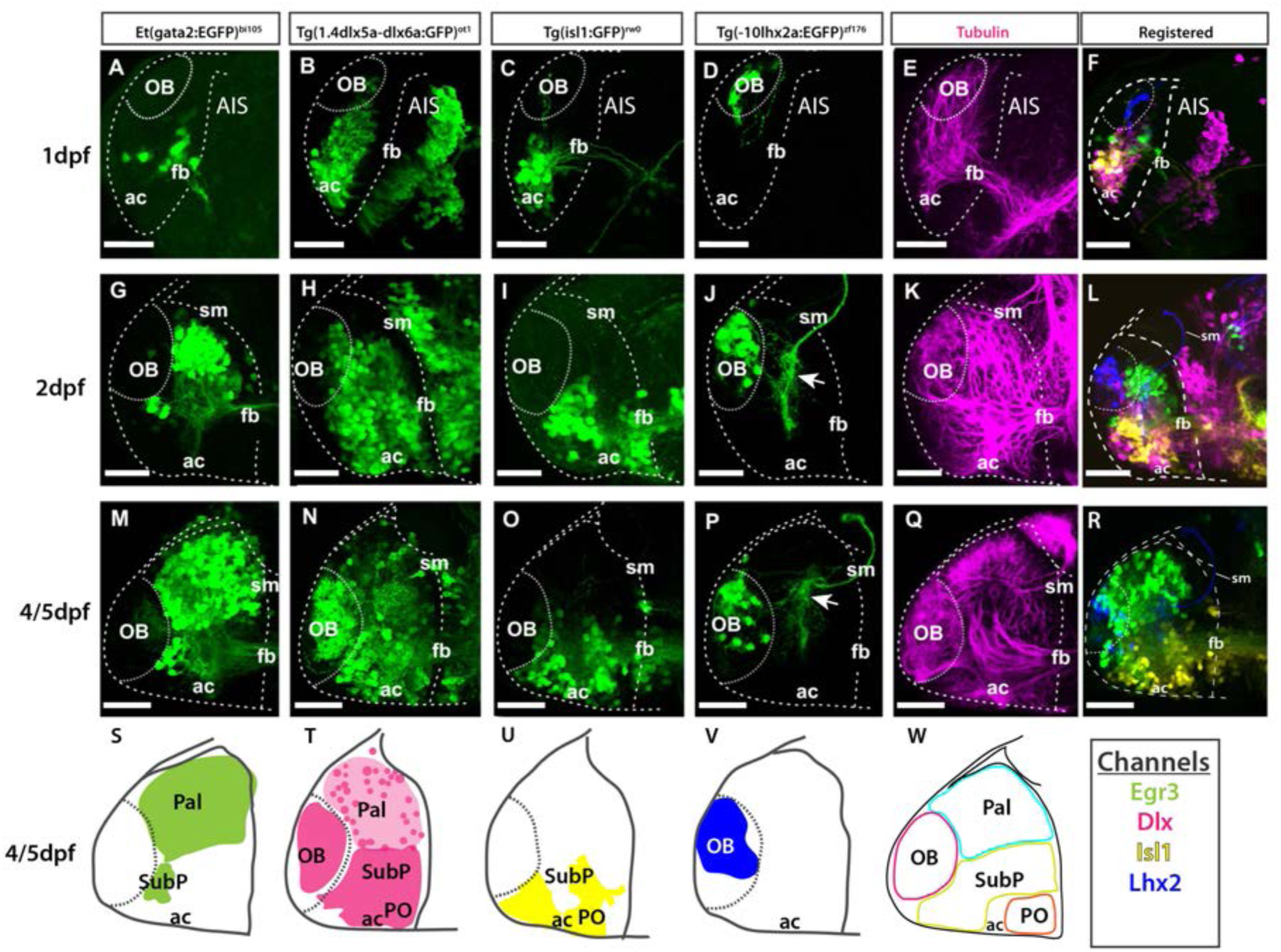
Atlas of transgene expression in the developing telencephalon: Lateral views. Et*(gata2:EGFP)* ^bi105^ in comparison with Tg(*1.4dlx5a-dlx6a*:GFP)^ot1^, Tg(*isl1*:GFP)^rw0^, Tg(-*10lhx2a:EGFP*)^zf176^ and acetylated tubulin antibody staining. Lateral views of 1dpf, 2dpf and 5dpf fish stained against GFP and acetylated tubulin **(E,K,Q).** Images are projections of confocal stacks. Rostral to the left. **M**: 4dpf; **N-R**: 5dpf. **S-W:** Schematics showing the location of the GFP+ cells in the different transgenic lines. **W:** Schematic showing all GFP+ domains. **F,L, R:** Transgenic datasets registered to a single reference brain for each developmental stage. Channels key below final column shows the colour of each transgenic in the registered images and the schematics. Scale bars: 50μm.

**Figure 8:**
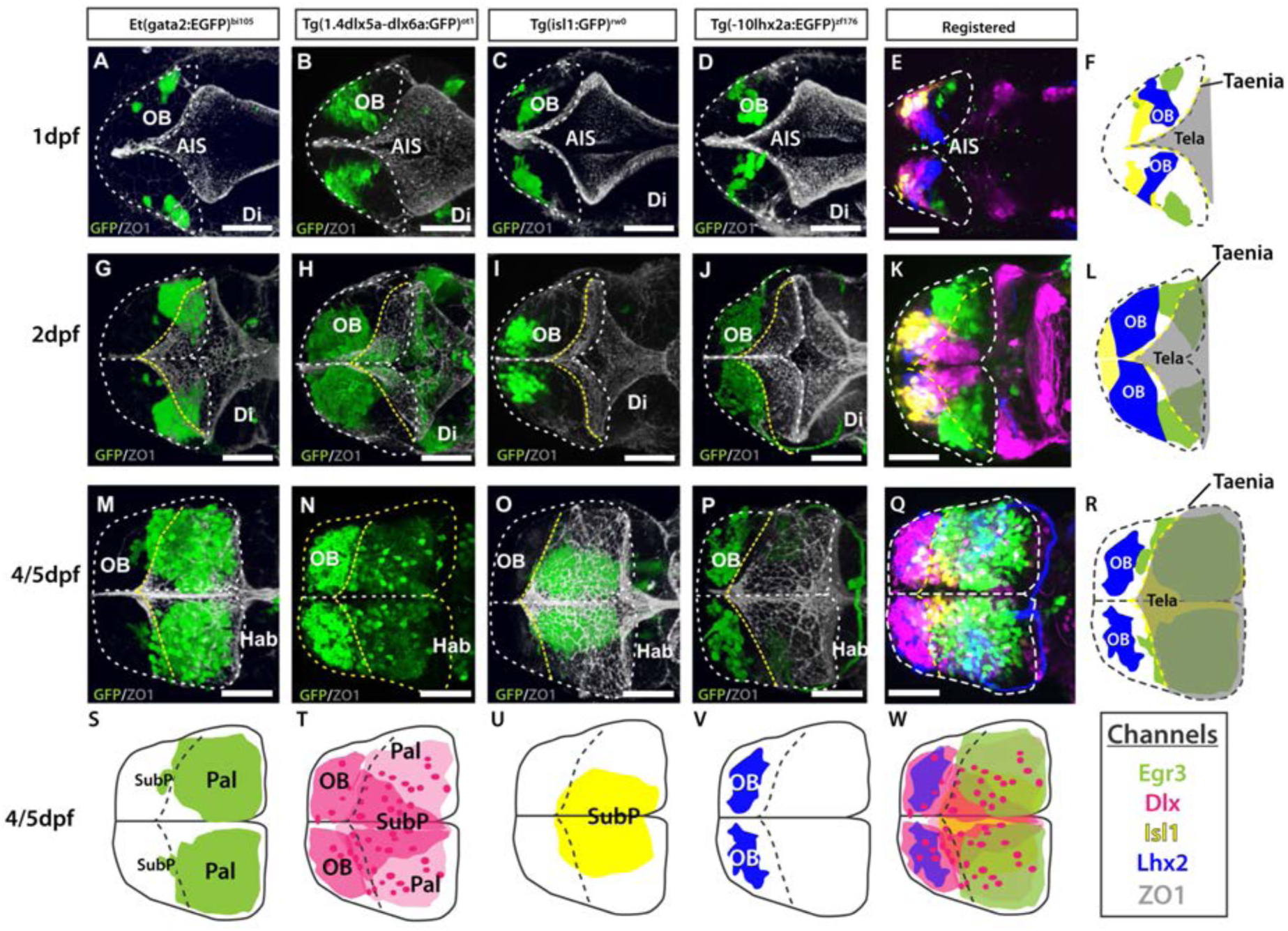
Atlas of transgene expression in the developing telencephalon. Dorsal views. Et*(gata2:EGFP)* ^bi105^ in comparison with Tg(*1.4dlx5a-dlx6a*:GFP)^ot1^, Tg(*isl1*:GFP)^rw0^, Tg(-*10lhx2a:EGFP*)^zf176^. ZO1: ventricular zone and *tela choroidea*. Dorsal views of 1dpf, 2dpf and 5dpf fish stained against GFP and ZO1. Images are projections of confocal stacks. Rostral to the left. **M**: 4dpf. **N-P**: 5dpf. White dashed line in A-R marks the outline of the telencephalon. Yellow dashed marks in A-R the rostral attachments of the *tela choroidea* or *taeniae.* **F, L & R:** Schematics showing the location of GFP+ cells in each transgenic line relative to the ZO1 positive *tela choroidea* (semi-opaque grey) over development, showing the expansion of the *tela* and rearrangement of telencephalic domains. The *taenia* is the point of attachment of the *tela choroidea* . **S-V**: Schematics showing the location of the GFP+ cells in each transgenic dataset by 4-5dpf. Black dashed line marks the rostral attachments of the *tela choroidea* or *taeniae*. **E, K, Q:** Transgenic datasets registered to a single reference brain for each developmental stage. **W:** Schematic of all registered GFP+ expression. Channels key below final column shows the colour of each transgenic within the composite images and schematics. Scale bars: 50μm.

#### Olfactory Bulb

To view changes in olfactory bulb position and cell populations during early telencephalon development, the following transgenes were used in combination with the Et*(gata2:EGFP)*^bi105^ transgene in registered data sets:

1. Tg(-*10lhx2a:EGFP*)^zf176^ (Fig. 7D, J, P, V; Fig. 8D, J, P, V) labels a subset of mitral cells, the principle output neurons of the olfactory bulb (Miyasaka et al., 2009, 2014).
2. Tg(*1.4dlx5a-dlx6a:GFP*)^ot1^ (Fig. 7B, H, N, T; Fig. 8B, H, N, T) labels a subset of olfactory bulb GABAergic interneurons (Li et al., 2005). Figure 9 shows Tg(*1.4dlx5a-dlx6a:GFP*)^ot1^ in comparison with Et*(gata2:EGFP)*^bi105^ in frontal view.
3. Anti-acetylated tubulin immunohistochemistry labels olfactory bulb glomeruli, amongst other structures (Fig. 7 E, K, Q; Fig. 9D, Fi; see Folgueira et al., 2012).

**Figure 9:**
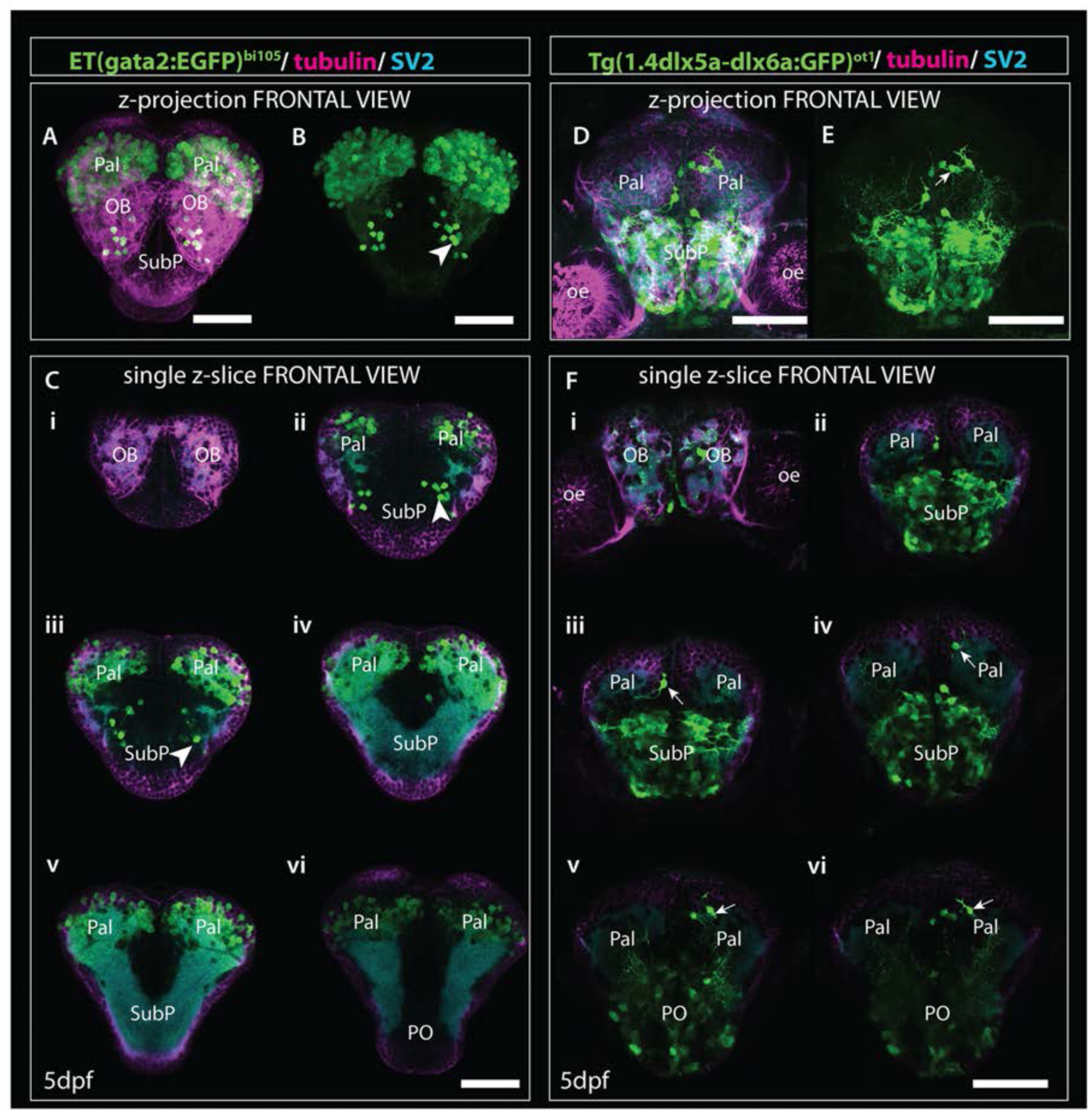
Comparison of gata2:EGFP and 1.4dlx5a-dlx6a:GFP expression in frontal view (z-projections and single z-slices). (A) Frontal view of the telencephalon of a 4dpf Et*(gata2:EGFP)*^bi105^ fish labelled with anti-EGFP (green), anti-tubulin (magenta) and anti-SV2 antibodies (cyan). **(B)** EGFP channel only. **(Ci-Cvi)** Single z-slices from rostral (i) to caudal levels (vi). (Arrowheads in B, Cii and Ciii) point to gata2:EGFP + subpallial neurons. **(D-E)** Frontal view of the telencephalon of a 4dpf Tg(*1.4dlx5a-dlx6a*:GFP)^ot1^ fish labelled with anti-EGFP (green), anti-tubulin (magenta) and anti-SV2 antibodies (cyan). **(E)** GFP channel only. **(Fi-Fvi)** Single z-slices from rostral (i) to caudal levels (vi). (Arrows in E and Fiii-Fvi) label *1.4dlx5a-dlx6a*:GFP+ interneurons in the pallium. Ci-Cvi and Fi-Fvi are equivalent rostrocaudal levels. Scale bars: 50μm.

The first mitral cells to differentiate by 1dpf (Figs. 7D, F, 8D, E) were located within the dorsal telencephalon, close to the anterior intra-telencephalic sulcus (AIS). By this stage, mitral cells had already extended processes (Fig. 7D). Between 2dpf (Figs. 7J, 8H) and 5dpf, the telencephalic domain caudal to the olfactory bulb expanded greatly, increasing the distance between the OB and the AIS (Figs. 7P, R, 8P, Q). Mitral cell axons reached the pallium and right habenula via the olfactory tracts and *stria medullaris* respectively (Fig. 7J, P, 8P-Q; see also Miyasaka et al., 2009, 2014). Olfactory bulb interneurons, visualised with the *1.4dlx5a-dlx6a:GFP* transgene, could be differentiated based on cell morphology. By 2dpf, these interneurons already extended processes towards the olfactory glomeruli (not shown), which increase in morphological complexity from 2dpf to 5dpf (Figs. 7N, 8N, 9Fi).

#### Pallium

The development of the pallium, as visualised through the changes in expression of *gata2:EGFP* in *Et(gata2:EGFP)*^bi105^ fish (Figs. 7A, G, M, S; 8A, G, M, S), has already been well covered in earlier parts of this paper. Here we compared EGFP expression with that in Tg(*1.4dlx5a-dlx6a:GFP*)^ot1^ fish, in which pallial GABAergic interneurons of subpallial origin are labelled (Mione et al, 2008; Yu et al., 2011), and in Tg(-*10lhx2a:EGFP*)^zf176^ fish. We observed that *1.4dlx5a-dlx6a:GFP*+ interneurons had a salt and pepper distribution in the pallium (Figs. 7N, 8N), being more sparse in number than gata2:EGFP*+* cells (Figs. 7M, 8M, Q, 9Fi-vi). Note that registration of Tg(*1.4dlx5a-dlx6a:GFP*)*^ot1^* data in lateral view (4dpf) in combination with the other markers was not satisfactory (hence not shown), but this was not the case for the dorsal view (Fig. 8Q) The morphological complexity of 1.4dlx5a-dlx6a:GFP+ pallial interneurons increased over time, such that by 4/5dpf they ramified dense processes throughout the pallial neuropil (Figs. 7N, 8N, Q, 9Fiii-iv). The -*10lhx2a:EGFP*^zf176^ transgene labels a neuropil within the pallium, within which axons from olfactory mitral cells terminate. This neuropil, which is nested within the gata2:EGFP+ pallial domain, may give rise to the posterior zone of the dorsal telencephalic area (labelled Dp in Fig S1 D-E; see Miyasaka et al., 2099;2014)

#### Subpallium

To observe how eversion affects subpallial populations, two transgenic lines were imaged in combination with the Et*(gata2:EGFP)*^bi105^ line.

1. Tg(*1.4dlx5a-dlx6a:GFP*)^ot1^ (Fig. 7B, H, N, T; 8B, H, N, T; 9D-F) labels neuronal precursors and GABAergic neurons in regions of the subpallium and preoptic region (MacDonald et al., 2010; Yu et al., 2011). Based on the expression pattern of *dlx5a* mRNA in the adult, this transgene is likely to label the dorsal (Vd), ventral (Vv), supracomissural (Vs), central (Vc), lateral (Vl) and posterior divisions of subpallium (Vp) (see Ganz et al., 2011). These divisions are not well differentiated by 5dpf, so we were unable to confirm this expression pattern.
2. Tg(*isl1:GFP*)^rw0^ (Fig. 7C, I, O, U & Fig. 8 C, I, O, U) (see Higashijima et al., 2000) shows a more restricted expression in the subpallium than the *1.4dlx5a-dlx6a:GFP*^ot1^ transgene. Recently, GFP expression in the adult brain has been extensively described in this transgenic line (see Baeuml et al., 2019) where the transgene labels neurons in the ventral subpallium (Vv), ventral part of the dorsal subpallium (Vdv) and ventral domain of the supracommisural subpallium (Vs)] (see Baeuml et al., 2019; Porter and Mueller, 2020). The transgene is also expressed in the preoptic region (see Baeuml et al., 2019).

At 1dpf, the ventral gata2:EGFP+ cluster overlapped with the *1.4dlx5a-dlx6a:GFP+* domain (Figs. 7A-B, F; 8A-B, E; 9A-B, Cii, D-E, Fii), but not with the isl1:GFP*+* one (Figs. 7C, F; 8C, E-F; see also S1). Pallial gata2:EGFP+ cells were located dorso-laterally to the 1.4dlx5a-dlx6a:GFP+ domain at this stage (Fig. 8A-B, E). At 2dpf, *1.4dlx5a-dlx6a:GFP* expression was largely complementary to *gata2:GFP* expression (Figs. 7G, H, L, 8G, H, K, L), marking the pallial-subpallial boundary (Fig. 8K). At this stage, viewing from a dorsal aspect (Fig. 8K) showed that the gata2:EGFP+ (pallium) and 1.4dlx5a-dlx6a:GFP*+* regions are distributed radially, with gata2:EGFP+ pallial cells located dorsolateral to the 1.4dlx5a-dlx6a:GFP*+* domain. This arrangement changes with elongation of the telencephalon along its rostro-caudal axis from 2dpf to 5dpf (Fig. 8M-Q, R-W).

#### Expansion of the tela choroidea

The *tela choroidea* is a thin sheet of neuroepithelial origin that encloses the dorsal part of telencephalic ventricle of the everted telencephalon. At 1dpf, the telencephalic ventricle, delineated by anti-ZO1 labelling (this antibody labels apical cell junction proteins in both the *tela choroidea* and the cells lining the ventricle, see Fig. S2), is restricted medially, becoming larger caudally at the level of the AIS (Fig. 8A-D, F). At this stage, both 10lhx2a:EGFP+ mitral cells (Fig. 8D, F) and gata2:EGFP+ pallial cells (Fig. 8A, F) were not covered by the ventricular surface and the *tela choroidea* dorsally, thus the telencephalon was not yet everted. By 2dpf, the number of gata2:EGFP*+* and 1.4dlx5a-dlx6a: GFP positive cells increased, with the caudalmost telencephalic parenchyma expanding into the AIS (Fig. 8G-H, K-L). Thus, a portion of the pallium becomes covered by the ventricular surface and associated *tela choroidea*, so from this stage the telencephalon (pallium) can be described as everted (Fig. 8G, L). By 5dpf most pallial cells are covered by the *tela choroidea* with the exception of very few gata2:EGFP*+* pallial cells located rostrolaterally (Fig. 8M, 8R). This shows that the pallium is not completely everted by 5dpf (Fig. 8M, R), but it will be at later stages. Thus, it seems that eversion process does continue after 5dpf, at least at extreme rostral locations. The olfactory bulb was clearly not everted, as the rostral attachment of the *tela choroidea* or *taenia* (for full details on the *taeniae* and eversion see Nieuwenhuys, 2009b) was just caudal to the olfactory bulb (Fig. 8P, V).

In addition to registering these transgenes at key stages during telencephalic eversion, we also imaged the Et*(gata2:EGFP)*^bi105^ transgenic line at 6dpf, the stage of development used in the Zebrafish Brain Browser atlas (ZBB). Figure S1 shows a snapshot of the registered Et*(gata2:EGFP)*^bi105^ expression pattern as viewed in the online ZBB viewer. A .PNG file is available for download (see supplementary data) so that users may upload this expression pattern and compare/combine it with any other labels registered to ZBB.

### 4.6. *gata2*:*EGFP* expression in 4dpf and 20dpf larval stages

Although the focus for this study has been the telencephalon, *gata2*:*EGFP* is expressed at other sites in the developing and mature CNS that will be described here for 4dpf and 20dpf larval zebrafish.

#### 4dpf zebrafish

To characterise *gata2*:*EGFP*^bi105^ expression, we used anti-GFP immunocytochemistry combined with anti-SV2 counterstaining to highlight anatomical landmarks (Turner et al., 2014). In the secondary prosencephalon, outside the telencephalon, we found EGFP expression in the lateral hypothalamus (Fig. 10 A,B). In the diencephalon, we observed a few neurons in the left-sided parapineal organ (Turner et al., 2016 and Fig. 9C), superficial pretectum, lateral and medial areas of the posterior tuberculum. In the midbrain, scattered EGFP+ cell bodies in the periventricular gray zone of the optic tectum extended dendrites into the superficial tectal neuropil (Fig. 10A,C). Based on location and morphology, these EGFP+ neurons could represent periventricular interneurons [Robles et al., 2011; type XIV cells of Meek and Schellart (1978)/ small periventricular cells of Vanegas et al. (1974)]. In the hindbrain, several EGFP+ cell bodies were present in the superior raphe, areas of the reticular formation (Fig. 10B-B’’) and caudally in areas of the medulla oblongata (Fig. 10C-C’’). In all regions, expression appeared to be predominantly in neurons rather than ventricular cells/glia.

**Figure 10:**
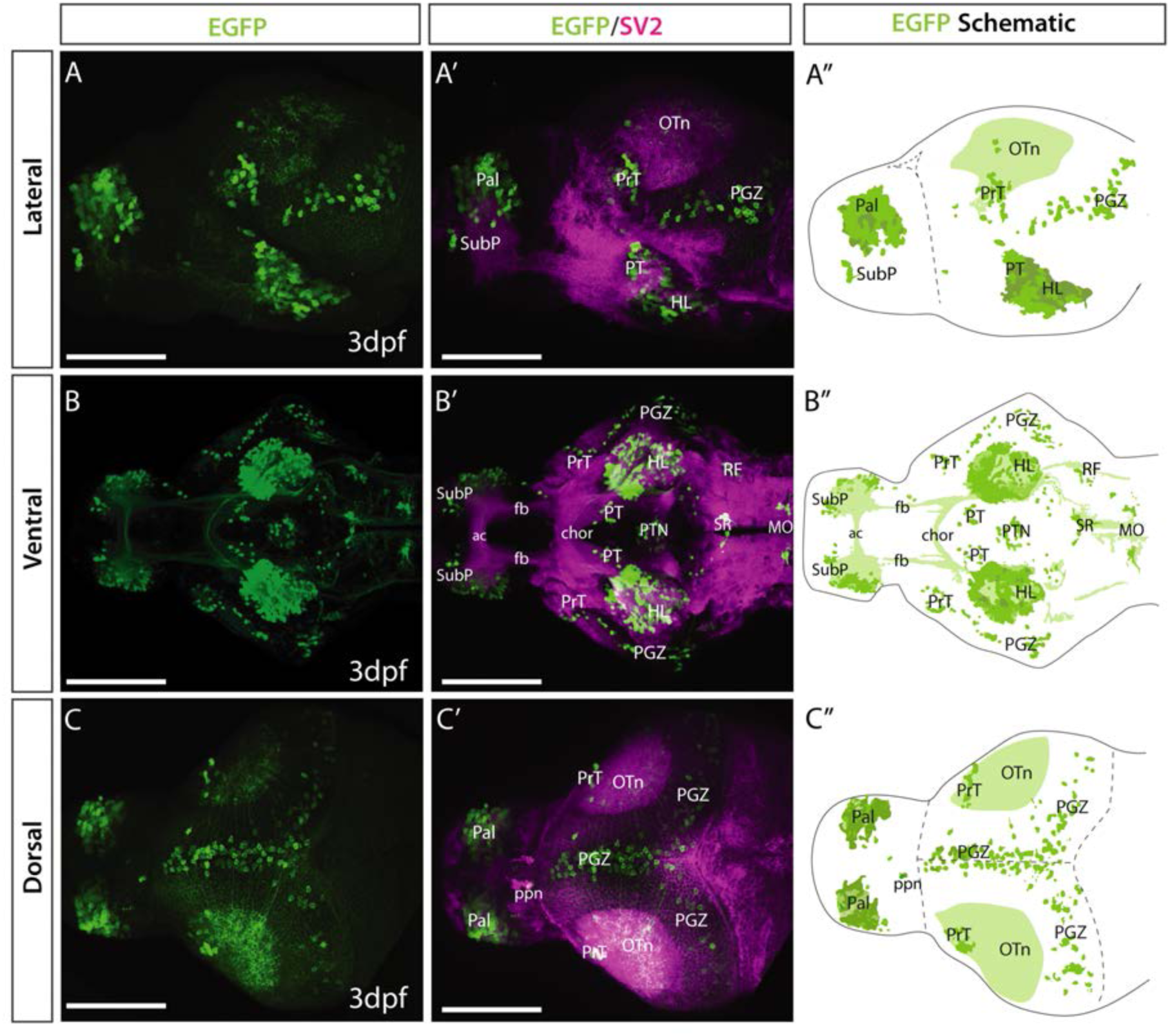
Et*(gata2:EGFP)*^bi105^ line EGFP expression at 3dpf. Lateral **(A-A’),** ventral **(B-B)** and dorsal **(C-C)** views of a 4dpf Et*(gata2:EGFP)*^bi105^ larvae labelled with anti-EGFP (green) and anti-SV2 antibodies (magenta). Major areas of EGFP expression are annotated in the schematics on the right (**A”, B” and C”**), these include: pallium, subpallial nucleus, pretectum, hypothalamic areas, optic tectum and hindbrain areas, among others. Scale bars: 100μm.

Despite the EGFP being a cytoplasmic variant, transgene expression (detected by immunohistochemistry) labelled several tracts and neuropil domains (Fig. 10B). The superficial tectal neuropil was EGFP+ (Fig. 10A-A’’, C-C’’); most of these fibres appeared to originate from cell bodies located in the periventricular gray zone, however some could have other origins. The horizontal commissure in the hypothalamus also contained EGFP+ fibres (Fig. 10B-B’’). Finally, at caudal rhombencephalic levels, EGFP+ fibres were present in the medial (MLF) and lateral (LLF) longitudinal fascicles.

#### 20dpf zebrafish

EGFP expression was more scattered at 20dpf (Fig. 4) than 4dpf (Fig. 8), but expression areas were consistent with those at 4dpf. At 20dpf, there was more mosaicism in expression within and between individuals, suggesting transgene silencing over time. This mosaicism was even more prevalent in adult transgenic fish (not shown). The only regions with *de novo* expression observed were in the cerebellum and vagal lobe (Fig. 4K).

In the basal secondary prosencephalon, we observed EGFP+ cells in the inferior hypothalamic lobes (Fig. 4G-J). In the alar diencephalon, we observed that the parapineal (ppn) was labelled, as previously described in Turner et al. (2016) at 1mpf (Fig. 4F and inset ii). EGFP+ fibers from the left-sided parapineal innervate the left dorsal habenula. At the same level as the habenulae, a small group of EGFP+ pretectal cells were present just above the forebrain bundle (Fig. 4F). In the basal diencephalon, EGFP+ cells were found in the preglomerular complex, torus lateralis, scattered cells in anterior and posterior tuberal regions (Fig. 4G-J).

In the midbrain and hindbrain there were only a few areas of expression. Transverse sections at the level of the optic tectum showed EGFP+ cell bodies in the PGZ and in the stratum album central of the optic tectum (Fig. 4G-I) that extended processes, most probably dendrites, dorsally into the stratum fibrosum griseum superficially and stratum opticum, consistent with 4dpf (see above). Also consistent with 4dpf, in the hindbrain, we observed EGFP+ neurons in the superior and intermediate reticular formation (Fig. 4J and inset iii) and superior raphe (Fig. 4K). In addition, we observed faintly EGFP+ cells in the valvula and corpus cerebelli (Fig. 4K and inset iv), likely to represent eurydendroid and/or Purkinje cells, and EGFP+ cells in the vagal lobe (Fig. 4L). These last two populations are additional to those observed in the 4dpf larva.

In summary, our analyses show EGFP expression in *Et*(*gata2*:*EGFP*)^bi105^ transgenic fish is restricted to specific areas in the brain that are broadly consistent between 4dpf and 20dpf.

## 5. Discussion

In this study, we have performed a detailed analysis of spatially restricted transgene expression in the telencephalon, in combination with other markers, to follow eversion and regionalisation of the zebrafish telencephalon during larval development. The registration of data from transgenic lines and other structural markers onto reference telencephali at key timepoints during development provides a framework atlas of major regions, structures and cell populations within the developing telencephalon. These datasets are available for further refinement through addition of further datasets and incorporation within other atlases, such as the Zebrafish Brain Browser (Marquart et al., 2015, 2017) within which we have registered a 6dpf Et*(gata2:EGFP)*^bi105^ dataset (Supp Fig. S1).

### 5.1. An atlas of telencephalon development

To understand how neural circuits control behaviour it is necessary to integrate well-annotated structural data with functional data (Arrenberg and Driever, 2013). This is challenging with respect to the teleost telencephalon that exhibits an everted morphology, rendering direct comparison with other vertebrate telencephali difficult. The repertoire of tools available for dissecting the anatomy and function of neural circuits has increased dramatically in the last few years (Scott, 2009; Leung et al., 2013; Feierstein et al., 2015; Föster et al., 2017, 2018; Robles et al., 2017), and includes the creation and characterisation of many transgenic lines expressing fluorescent proteins in discrete brain areas (Kwan et al., 2007; Scott et al., 2007; Takeuchi et al., 2015; Kawakami et al., 2010; Suster et al., 2011; Randlett et al., 2015; Marquart et al., 2017). Despite this wealth of tools and resources, there is still need of well-annotated baseline structural data that can be used to annotate reference brains (Ronnenberger et al., 2012; Marquart et al., 2015, 2017; Randlett et al., 2015).

Existing neuroanatomical resources for zebrafish include 3D atlases based on confocal data during development and in the adult (Bryson-Richardson et al., 2007; Ullmann et al., 2010; Ronnenberger et al., 2012; Marquart et al., 2015, 2017; Randlett et al., 2015; Kunst et al., 2019; Kenney et al., 2021; zebrafishbrain.org). However, these atlases are usually based on a single developmental timepoint, so cannot communicate the gross morphological changes happening early in development. Here we present an atlas of the zebrafish telencephalon during key stages of development (1dpf, 2dpf and 5dpf). Our 5dpf annotations are easily extended to 6dpf, the stages presented in the Zebrafish Brain Browser (Marquart et al., 2015, 2017), Zebrafish Brain Atlas (Randlett et al., 2015) and mapzebrain (Kunst et al., 2019).

We have used anti-acetylated tubulin as an anatomical reference instead of cytoplasmic/nuclear staining often used in other atlases (Ronnenberger et al., 2012; Marquart et al., 2015; Radlet et al., 2015). We observed that, as previously shown in VibeZ (Ronnenberger et al., 2012), this labelling works well as a reference channel and has the added bonus that one can view the development of the axonal scaffold through development. We also include the marker ZO1, which delineates the extension of the ventricular zone and allows the degree of eversion to be tracked. Given there are portions of the ventricle that, as a result of eversion, are difficult to visualize without markers (Nieuwenhuys, 2009b), ZO1 or other ventricular zone markers are crucial when building a reference telencephalon resource.

### 5.2. Eversion and “outside-in” construction mode of the teleost telencephalon

Eversion models have failed so far to explain how morphogenetic events during development lead to the organization of the pallial regions in the adult (Yamamoto et al., 2007; Mueller et al., 2011; Porter and Mueller, 2020). After the primary events of eversion from 1dpf to 5dpf (Folgueira et al., 2012), there is a subsequent major expansion of the telencephalon that affects its organization (present results; Dirian et al., 2014; Furlan et al., 2017). With the main events of eversion occurring so early (mostly by 1-5dpf) and growth so late, this complicates the eversion narrative of a simple lateral out-folding of the neural tube during early development (Butler and Hodos, 2005; Yamamoto et al., 2007; Nieuwenhuys, 2011; Mueller et al., 2011; Porter and Mueller, 2020). For instance, eversion models currently do not adequately account for differential timing in the development of different telencephalic areas.

The relative size of different telencephalic domains changes dramatically over the first weeks of development. For instance, early on, the nascent olfactory bulbs constitute much of the telencephalon whereas there is disproportionate, massive expansion of the pallial telencephalic lobes from 5dpf to 20dpf. Within the expanding pallium, Dirian et al. (2014) found two distinct populations of neural stem cells that are segregated in space: a dorso-medial domain and a lateral domain. Consistent with late expansion of pallial territories, the lateral domain that contributes neurons to the lateral domain of the pallium, Dl, only becomes neurogenic from 5dpf onwards.

Following morphogenesis of the ventricle, tangential growth of the ventricular zone, driven by telencephalic neuronal differentiation, may be one of the driving forces to expand the pallial ventricular surface (Aboitiz and Montiel, 2019; present results). By 5dpf, the zebrafish telencephalon shows everted morphology (Folgueira et al., 2012; present results), but there is a subsequent expansion of the telencephalic ventricular zone, which grows in lockstep with general telencephalic lobe growth. Cells are added in an outside-in manner following a “sequential stacking” mode, with neurons arranged in age-related layers surrounding a central “core” of earliest-born neurons (Furlan et al., 2017). This addition of newborn cells towards the pallial ventricular surface may force tangential expansion of the ventricular zone as the telencephalon grows. We have observed that EGFP expression in Et*(gata2:EGFP*)^bi105^ larvae appears to represent post-mitotic telencephalic cells, thus, the first cells to express EGFP in this line probably constitute this central “core”.

### 5.3. gata2:EGFP^bi105^ + pallial neuron morphologies

Using a Crispr-based labelling technique, we revealed morphologies of cell types in the pallium and subpallium of 6dpf larvae. We observed two different cell types in the pallium by 6dpf that, despite the early age of the animals, show quite elaborate spiny dendritic arbours, similar to those reported in adult fish. Studies in various adult teleosts have shown different neuronal cell types occupying different areas of the pallium (Demski and Beaver, 2001; Demski, 2013). Deep pallial territories host large efferent projection neurons, while periventricular neurons are predominantly small stellate shaped cells (Demski and Beaver, 2001; Demski, 2013). These findings match well with the cell types described in this study. Stellate cells reported here, for instance, resemble the morphology of interneurons present in lateral and medial regions (Dm) of the pallium in adult fish (Yamane et al., 1996; Demski and Beaver, 2001; Giassi et al., 2012; see Demski, 2013 for a review). Thus, even at early stages, differentiated pallial neurons already show complex arborisation, indicative of being integrated in active neural circuits.

### 5.4. Establishment of neural circuits and the emergence of control of behaviours in the telencephalon

The adult telencephalon is involved in many functions, such as motor control, sensory processing, memory, learning, emotions and social interaction. What is the sequence of maturation of the telencephalic circuits involved in these functions? Our results indicate differentiation of neurons in the olfactory bulb first, while areas of the pallium and subpallium show considerable expansion after 5dpf suggesting late maturation of these regions. This is in agreement with behavioural studies, which have shown that complex behaviours appear from 2 to 6 weeks post-fertilization (Valente et al., 2012; Aoki et al., 2013; Cheng et al., 2014; Dreosti et al., 2015; Lal et al., 2018; Stednitz et al., 2018; Tunbak et al., 2020; Bartoszek et al., 2021). As a consequence, most functional studies likely involving the role of the telencephalon, such as fear conditioning, social interaction, learning and memory, performed on older zebrafish.

Despite late maturation of some telencephalic circuits, the animal needs to process information from the environment and coordinate movement from an early age. By 4dpf, the larva has already hatched and is transitioning into independent feeding. In this sense, we observed quite mature cell morphologies already by 2dpf in the olfactory bulb, indicative of early olfactory processing. We also observed that telencephalic gata2:EGFP+ cells show complex morphologies and connections by 5-6dpf. In the case of the subpallial gata2:EGFP+ cells, by 6dpf they already show a complex connection patter with caudal regions, which resembles the connections described in the adult for the subpallium (Rink and Wullimann, 2004). Thus, from an early age, this circuit might allow the young animal to exert telencephalic control over certain behaviours.

### 5.5. gata2:EGFP+ subpallial cells may be part of the zebrafish septum

Various cell populations and nuclei in the zebrafish subpallium have been described based on topographical location and neurochemistry. Among these, the ventral area (Vv) and the lateral area (Vl) of the subpallium are proposed to be homologous to septal areas in tetrapods (Rink and Wullimann, 2004; Wullimann and Mueller, 2004b; Ganz et al., 2011; Baeuml et al, 2019). Here we were able to follow the development of a gata2:EGFP+ subpallial cluster located at pre-commisural and commissural levels of the zebrafish telencephalon and analyse its connections. We identify gata2:EGFP+ subpallial cells with elaborate dendritic arbors that span around the anterior commissure and axons that bifurcate and project unilaterally or bilaterally to tuberal and hypothalamic areas. These cell populations could correspond, at least partially, with early-migrated telencephalic area M4 (see Mueller and Wullimann, 2005; Mueller et al., 2008). Previously we showed the prethalamic origin of another group subpallial cells in the zebrafish pallium (see Turner et al., 2016). On the contrary, we did not analyse cell migrations into the telencephalon in the present study, so we cannot comment on the exact origin of the subpallial gata2:EGFP+ cells, or, for that matter, any other population in the telencephalon.

Based on location and projection pattern, we believe gata2:EGFP+ subpallial cells could correspond to septal cholinergic populations described in the lateral subpallium of adult zebrafish (Mueller et al., 2004) and other teleosts (trout: Pérez et al., 2000). Both subpallial gata2:EGFP+ cells and cells immunoreactive for choline acetyl transferase (ChAT) share similar location in Vl and similar projection pattern to the hypothalamus (Mueller and Wullimann, 2004; Rink and Wullimann, 2004). Cholinergic cells also show varicose fibers in the subpallium, especially dense around the anterior commissure (Mueller and Wullimann, 2004), similar to the gata2:EGFP+ subpallial cells described in this study. We do not think that the location of the subpallial gata2:EGFP*+* cells correspond with neuropeptide Y+ cells previously described in Vl (Castro et al., 2006; Turner et al., 2016), which project to the dorsomedial pallium (not shown), or with the proper entopeduncular nucleus (former ventral entopeduncular nucleus; Turner et al., 2016). In any case, we have not looked at colocalization of either neuropeptide Y or ChAT in Et(*gata2:EGFP*)^bi105^ fish, so the putative identity of these gata2:EGFP*+* subpallial neurons remains speculative. The heterogeneity of cell types in the lateral subpallium indicates that further detailed analysis is needed to dissect nucleus identity, anatomy, connectivity and development of this area.

### 5.6. *egr3* expression and function, future directions for looking at telencephalic activity

The insertion point in Et(*gata2:EGFP*)^bi105^ fish is 7kb upstream of the first exon of *Early Growth Response 3 gene* (*egr3)*. Although so far we have used this transgenic line for neuroanatomical studies only (Folgueira et al., 2012; Turner et al., 2016; present study), we believe it will be a useful tool for toher analyses. Although the line expresses a stable form of EGFP, the CRISPR–Cas9 switching method (Auer et al., 2014; Kimura et al., 2014) could be used to convert this EGFP transgene to Gal4 to generate a more versatile transgenic line for such studies. Our results, using this method to interrogate single cell morphology, already show that the eGFP sgRNAs can effectively lead to transgene replacement.

*egr3* is a member of the immediate early gene (IEG) family that encodes transcription factors with almost identical zinc finger DNA binding domains (O’Donovan et al., 1999). Among other functions, IEGs such as *egr1, egr2/krox20, egr3* and *egr4* have been implicated in neural plasticity in response to neuronal activation (Li et al., 2005, 2007; Kim et al., 2012). In mice, *egr3* expression is induced by synaptic activity and is required for hippocampal long-term potentiation and long-term depression (Gallitano-Mendel et al., 2007), as well as for hippocampal and amygdala dependant learning and memory (Li et al., 2007). The persistent expression of *egr3* in spatially restricted populations of CNS cells from early stages (this study, Deguchi et al., 2009) suggests that in addition to roles in synaptic plasticity, the gene may also have a role in early neuronal differentiation or function. Indeed, a closely related gene *egr2/krox20,* is required for normal rhombomere development (Mechta-Grigoriou et al., 2000).

## 6. Abbreviations

ac: anterior commissure
AIS: anterior intra-telencephalic sulcus
CC: corpus cerebelli;
chor: horizontal commissure
Di: diencephalon
E: eye
EG: eminentia granularis
EmT: eminentia thalami
fb: forebrain bundle
GC: granularis cerebelli
Hb: habenula
HL: lateral hypothalamic lobe
Hyp: hypothalamus
lHb: left habenula
lfb: lateral forebrain bundle
LLF: lateral longitudinal fascicle
lot: lateral olfactory tract
LX: vagal lobe
mlf: medial longitudinal fascicle
MO: medulla oblongata
MR: median raphe
OB: olfactory bulb
Och: optic chiasm
Oe: olfactory epithelium
OT: optic tectum
OTn: optic tectum neuropil
P: pineal organ
Pal: pallium
PG: preglomerular complex
PGZ: periventricular grey zone
PHL: posterior hypothalamic lobe
PO: preoptic region
ppn: parapineal organ
PrT: pretectum
PT: posterior tuberculum
PTh: prethalamus
PTN: posterior tuberal nucleus
rHb: right habenula
RF: reticular formation
sm: stria medullaris
SR: superior raphe
SubP: subpallium
Tel: telencephalic lobes
TL: torus longitudinalis
TLa: torus lateralis hypothalami
vENT: ventral entopeduncular nucleus
Vv: ventral area of the subpallium
VIII: 8^th^ cranial nerve root

## 9. Author Contributions

MF, KJT and TAH conceived and designed the work, with inputs from SWW. MF, KJT, and TAH acquired and analysed the data. PH registered transgenic lines. KT, TAH and LEV developed the single cell labelling with Crispr/Cas9 technique. All authors contributed to interpretation of data and to writing the article. IHB provided comments on the final manuscript.

## 12. Acknowledgments

We thank members of UCL Fish Facility for help maintaining fish lines, members of our labs in UCL and UDC for discussion and support, Dr. Ramón Anadón for comments on initial draft of the manuscript, Dr. Thomas Auer for his advice and sharing of reagents for the single cell labelling using Crispr, and Dr. Tom Becker for the *Et(gata2:GFP)^bi105^* line.

## 13. Data Availability Statement

Authors are keen for their registered transgenic line data to be used as a tool for other researchers studying the developing telencephalon. Thus, authors will to make all raw data pertaining to this study available upon request.

---

**Figure S1:**
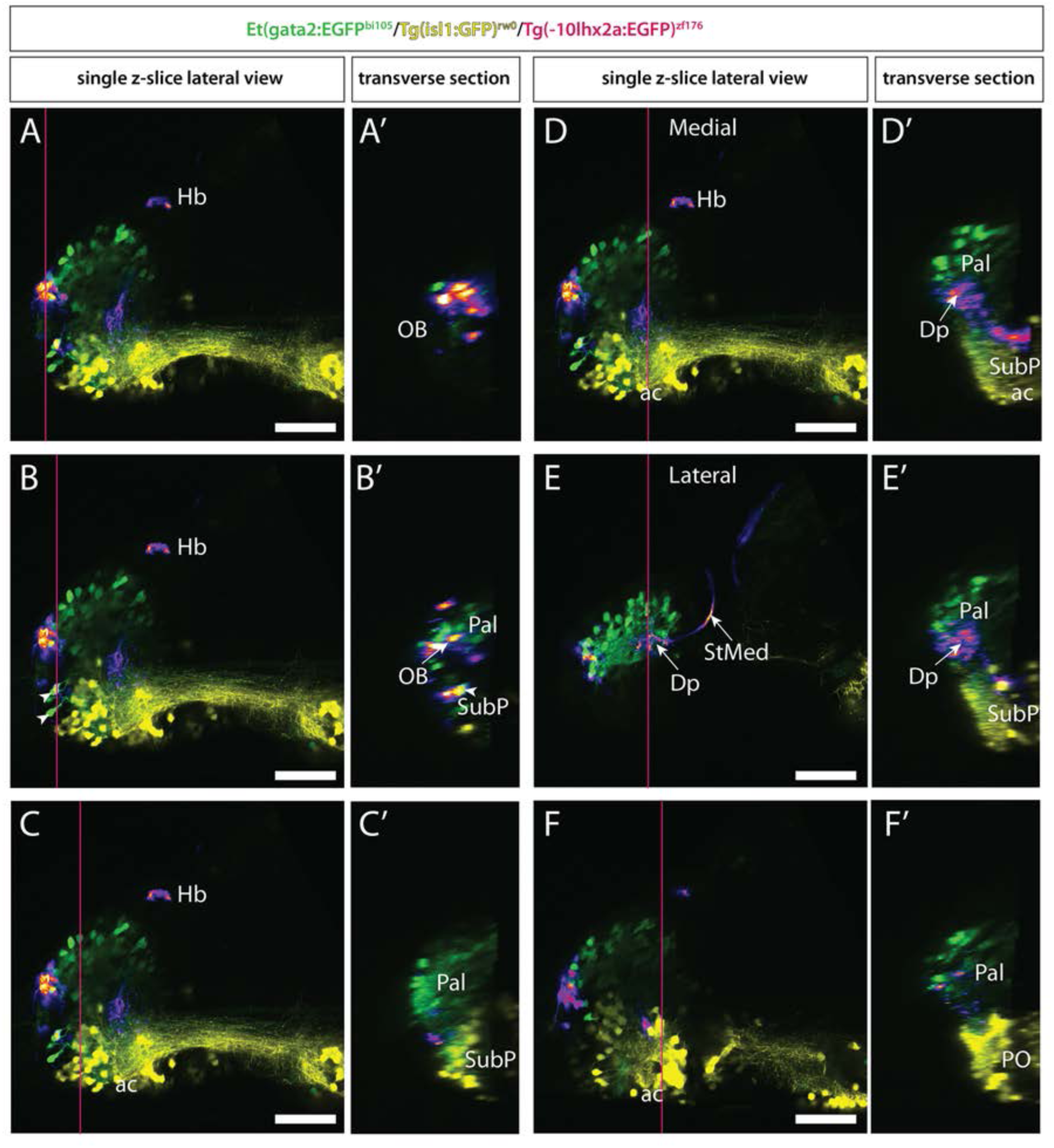
Registered brain (lateral view) showing Et*(gata2:EGFP)* ^bi105^ (green) , Tg(-*10lhx2a:EGFP*)^zf176^ (FIRE) and Tg(*isl1*:GFP)^rw0^ (yellow) transgene expression at 5dpf. A-F: Single z-slices showing transverse sections **(A’-F’)** made using orthogonal view tool (YZ) in Image J. E is more lateral than F. The rostro-caudal level of transverse sections **(A’-F’)** are indicated by a pink line on accompanying lateral z-slice in A-F. **D-E**: same rostro-caudal level, **D** closer z-slice to the midline than **E. D’-E’**: show the location of Tg(- *10lhx2a:EGFP*)^zf176^ mitral cell processes in putative Dp. Scale bars: 50μm.

**Figure S2:**
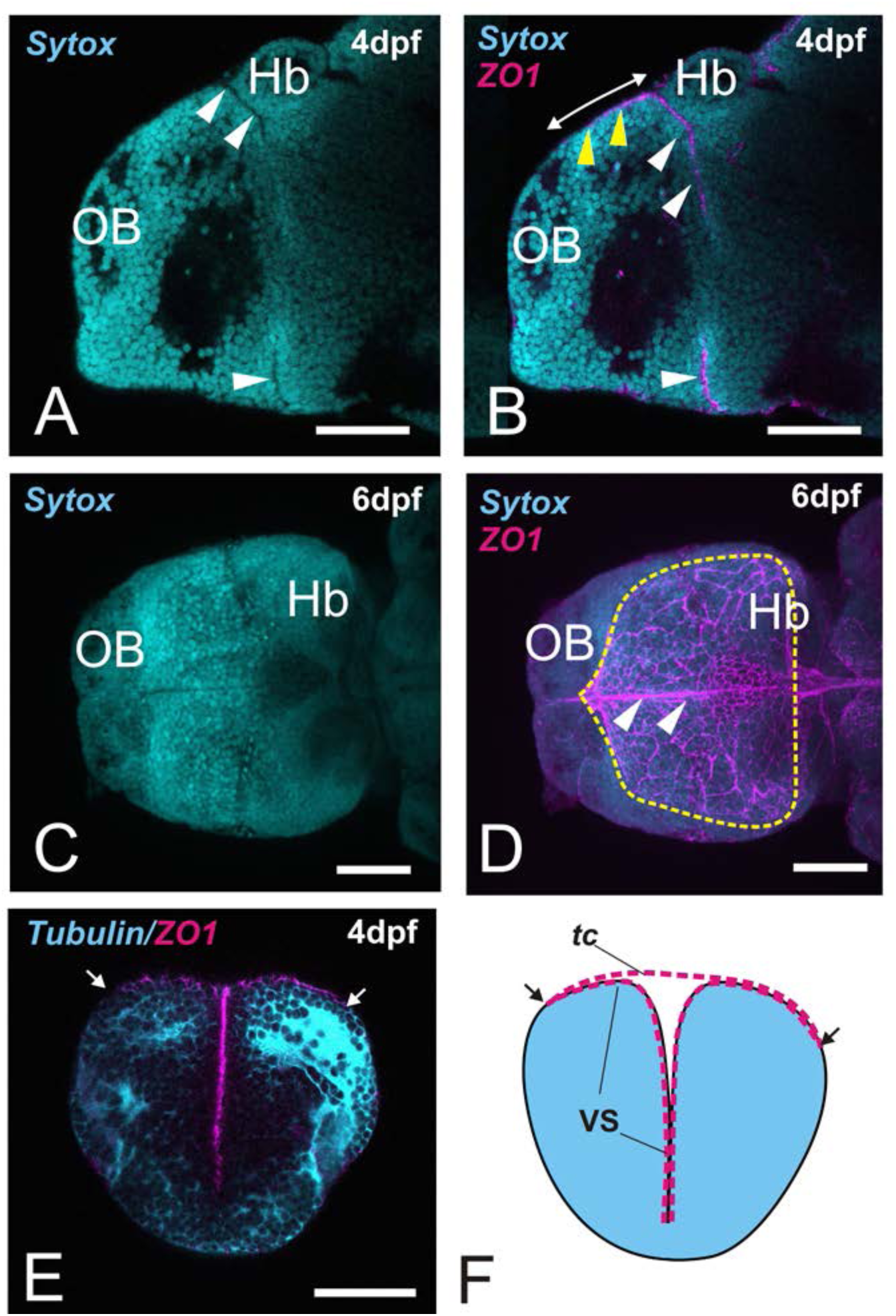
ZO1 as a marker of the ventricular surface and *tela choroidea*. Lateral (A-B) and dorsal views (C-D) of the telencephalon of 4dpf (A-B) and 6dpf (C-D) fish stained against ZO1 (magenta) and counterstained with sytox orange (cyan). **A-B:** White arrowheads point to the portions of the ventricle that are clearly visible with a nuclear stain only (AIS dorsally, optic recess ventrally). **B:** Yellow arrowheads point to the extension of the ventricle visible by ZO1 staining. Double arrow marks the extension of the *tela choroidea*. This could be easily missed with a nuclear staining only (compare A and B). **C-D:** Full extension of the ventricle and *tela choroidea* is not clearly visible with a nuclear stain only (C), but clear when ZO1 staining is used (dotted line marks the full extension of the ventricle and *tela choroidea* dorsally). **E**: Transverse section of the telencephalon of a 4dpf larva labelled against ZO1 (magenta) and tubulin (cyan). Note the T-shape of the ventricle. The *tela choroidea* extends dorsally (places of attachment or *taeniae* are marked with arrows). **F:** Schematic representation of E, showing the ventricular surface (VS) and *tela choroidea* (tc) labelled by ZO1. Brain parenchyma is represented in pale blue. Scale bars: 100.μm

**Figure S3:**
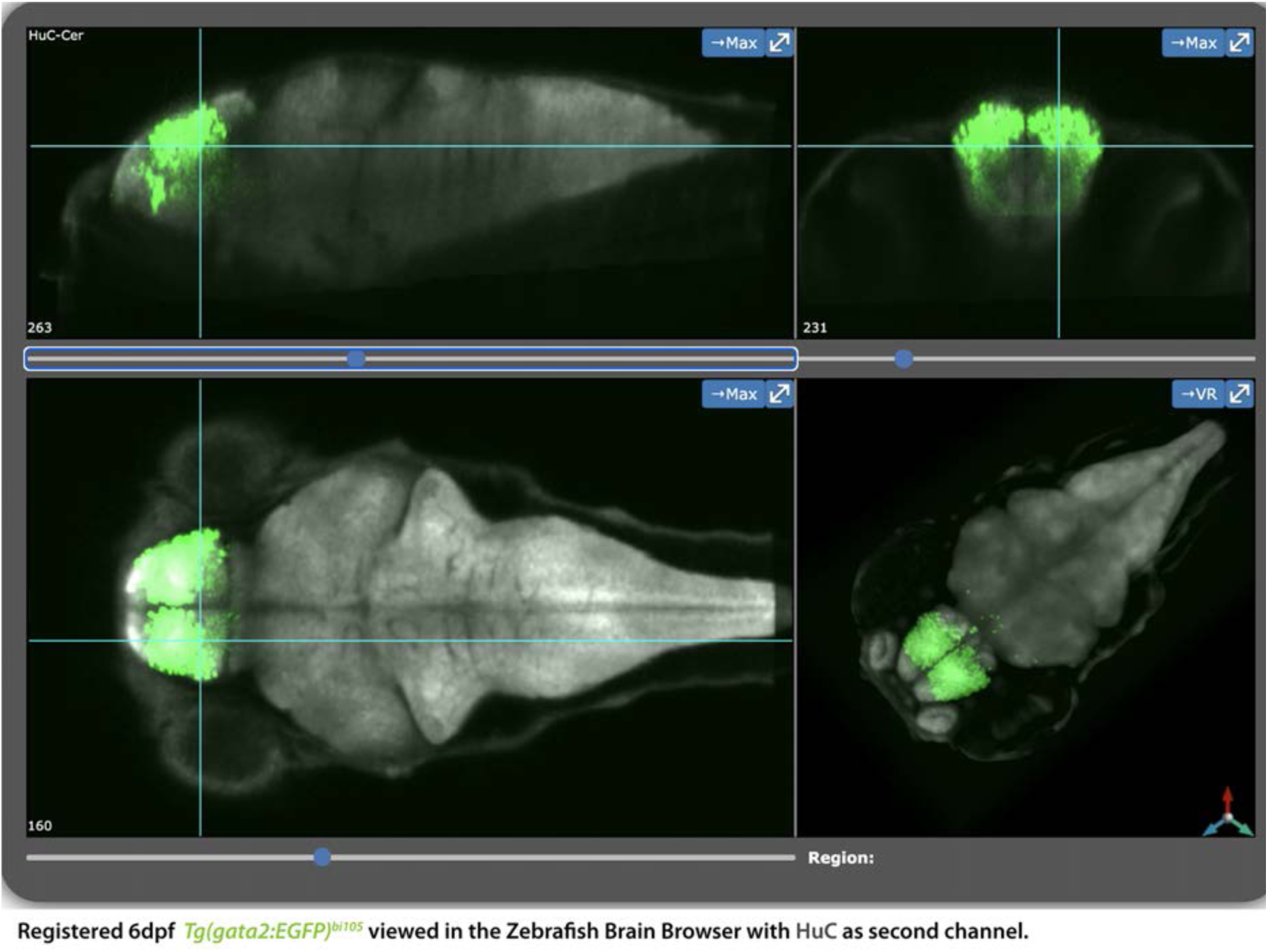
Et*(gata2:EGFP)*^bi105^ expression at 6dpf registered to Zebrafish Brain Browser (ZBB). Screenshot of the online ZBB viewer showing a 6dpf *Et(gata2:EGFP)*^bi105^ larvae labelled with anti-EGFP (green) registered to the ZBB standard brain. HuC channel (grey) shows full brain structure.

**Figure S4:** A 6dpf Et(gata2:EGFP)bi105 stack in PNG format. This dataset can be uploaded to Zebrafish Brain Browser atlas (see “Material and Methods”).

